# Circulating Lysophosphatidylcholines, Phosphatidylcholines, Ceramides, and Sphingomyelins and Ovarian Cancer Risk: a 23-year Prospective Study

**DOI:** 10.1101/565044

**Authors:** Oana A. Zeleznik, Clary B. Clish, Peter Kraft, Julian Avila-Pancheco, A. Heather Eliassen, Shelley S. Tworoger

## Abstract

**Background:** Experimental evidence supports a role of lipid dysregulation in ovarian cancer progression and metastasis. We estimated associations with ovarian cancer risk for circulating levels of four lipid groups measured 3-23 years before diagnosis.

**Methods:** Analyses were conducted among cases (N = 252) and matched controls (N = 252) from the Nurses’ Health Studies. We used logistic regression adjusting for risk factors to investigate associations of lysophosphatidylcholines (LPC), phosphatidylcholines (PC), ceramides (CER), and sphingomyelins (SM) with ovarian cancer risk overall and by histotype. A Bonferroni adjusted p-value threshold of 0.0125 (0.05/4; 4 measured lipid groups) was used to evaluate statistical significance. Odds ratios (OR; 10^th^ to the 90^th^ percentile) and 95% confidence intervals of ovarian cancer risk were estimated.

**Results:** C16:0 SM, C18:0 SM, C16:0 CER and SM sum were significantly positively associated with ovarian cancer risk, with ORs ranging from 1.95-2.10, with stronger ORs for postmenopausal women (2.02-3.22). ORs were generally similar for serous/poorly differentiated and endometrioid/clear cell tumors, although most did not meet the Bonferroni-adjusted p-value for significance. C18:1 LPC and the ratio of LPC to PC were significantly inversely, while C18:0 SM was significantly positively, associated with risk of endometrioid/clear cell tumors.

**Conclusion:** Elevated levels of circulating SMs 3-23 years before diagnosis were associated with increased risk of ovarian cancer, regardless of histotype, with stronger associations among postmenopausal women. Prospective and experimental studies are required to validate our findings and understand the role of lipid dysregulation, SMs in particular, in ovarian carcinogenesis.

## Introduction

Ovarian cancer is the fifth leading cause of cancer death for U.S. women with 4 out of 5 ovarian cancer patients diagnosed with advanced disease that has spread throughout the abdominal cavity (1). Identifying new risk factors for ovarian cancer is important to improve risk prediction models and discover new opportunities for prevention. One area of interest is lipid metabolism (2).

Laboratory evidence supports a role of lipid dysregulation in ovarian cancer progression and metastasis (3, 4). Several prospective human studies reported suggestive associations with complex lipids such as total cholesterol (positive) (5) or HDL (inverse) (6). However, there have been few comprehensive prospective studies of other lipids, such as lysophosphatidylcholines (LPC), phosphatidylcholines (PC), ceramides (CER) and sphingomyelins (SM), which appear to be different in ovarian cancer cases compared to healthy women (7-9).

LPCs act as signaling molecules involved in up-regulating cell proliferation, angiogenesis, migration, inflammation, and wound healing (3, 4, 10-15). Phospholipase A2 converts PCs to LPCs and may influence cell proliferation, invasion and migration (16). In multiple case-control studies, ovarian cancer cases had higher plasma or urine LPC levels compared to controls (17-26). However, due to the retrospective design of these studies, it is unclear whether alterations in these biomarkers preceded or followed the appearance of ovarian cancer.

Another class of lipids that may play a role in ovarian carcinogenesis are SMs that have a phosphocholine headgroup and ceramide (CER) backbone (16). CER is a pro-apoptotic signaling molecule in oocytes and ovarian tumors (27-30) and has potential metastasis suppressing properties (31). The primary source of ceramide in ovarian tumors is via SM metabolism. Ceramide levels are very low in ovarian tumors (27, 30, 32) while SMs appear to be higher in ovarian tumors versus normal tissue (33).

We leveraged novel metabolomics assays to measure four circulating lipid groups which have been previously hypothesized to be associated with ovarian cancer, LPCs, PCs, CERs and SMs, in plasma samples in a study of ovarian cancer risk within two large prospective cohorts with twenty-three years of follow-up.

## Methods

### Study Population

This analysis was based on data from nested case-control studies in the Nurses Health Studies (NHS (34) and NHSII (35). The NHS was established in 1976 among 121,700 US female nurses aged 30–55 years, and NHSII was established in 1989 among 116,429 female nurses aged 25–42 years. Participants have been followed biennially by questionnaire to update information on exposure status and disease diagnoses. In 1989–1990, 32,826 NHS participants provided blood samples and completed a short questionnaire (34). Details regarding blood collection (35) and identification of ovarian cancer cases are provided in the supplementary materials.

Cases were diagnosed with ovarian cancer three years after blood draw until June 1, 2012 (NHS), or June 1, 2013 (NHSII). Participants diagnosed with invasive disease and who died within 3 years of diagnosis were defined as rapidly fatal cases. Two hundred fifty-three cases of invasive and borderline epithelial ovarian cancer (213 in NHS and 40 in NHSII) were confirmed by medical record review. Cases were matched to one control on: cohort (NHS, NHSII); menopausal status and hormone therapy use at blood draw (premenopausal, postmenopausal and hormone therapy use, postmenopausal and no hormone therapy use, missing/ unknown); menopausal status at diagnosis (premenopausal, postmenopausal, or unknown); age (±1 year), date of blood collection (± 1 month); time of day of blood draw (±2 hours); fasting status (>8 hours or ≤8 hours). The study protocol was approved by the institutional review boards of the Brigham and Women’s Hospital and Harvard T.H. Chan School of Public Health, and those of participating registries as required.

### Metabolite profiling

Plasma metabolites were profiled at the Broad Institute of MIT and Harvard (Cambridge, MA) using a liquid chromatography tandem mass spectrometry (LC-MS) method designed to measure polar and nonpolar lipids as described previously (36-39). Details are provided in the supplementary materials.

Fifty-four metabolites were profiled in total. Metabolites with a coefficient of variation (CV) among the blinded QC samples higher than 25%, an intraclass correlation coefficient (ICC) <0.75, or with missing values in more than 10% of the participant samples were excluded from this analysis (N=3). Missing values in metabolites with less than 10% missing across participant samples were imputed with 1/2 of the minimum value measured for that metabolite. Metabolites not passing our previously conducted processing delay (N=12) pilot study (39) were also excluded from this analysis. All metabolites included in the analysis exhibited good reproducibility within person over one year (39).

Thirty-nine individual metabolites belonging to four metabolite classes (11 LPC, 17 PC, 6 SM and 5 CER) together with the sum of the measured metabolite values within each class (LPC sum, PC sum, SM sum and CER sum) and the ratios of LPC to PC (LPC:PC) and SM to CER (SM:CER) were analyzed in this study.

Metabolite values were transformed to probit scores for all subsequent analyses to reduce the influence of skewed distributions and heavy tails on the results and to scale the measured metabolite values to the same range.

### Statistical analysis

Conditional logistic regression was used to evaluate metabolite associations with risk of overall ovarian cancer among all participants, and separately by menopausal status at blood draw. Metabolite values were used as continuous variables to calculate linear trend p-values. We estimated the odds ratios (OR) and 95% confidence intervals (95% CI) for an increase from the 10^th^ to 90^th^ percentile in metabolite levels. Furthermore, we estimated OR of ovarian cancer and 95% CI for the SM sum measure across quartiles (based on the distribution in controls).

In a sensitivity analysis, we compared conditional logistic regression to unconditional logistic regression adjusting for the matching factors and found similar results (data not shown). Based on this comparison, subsequent analyses by histotype, rapidly fatal status and time between blood collection and diagnosis were conducted using unconditional logistic regression adjusting for the matching factors, allowing the use of all controls. All regression analyses were further adjusted for established ovarian cancer risk factors: duration of oral contraceptive use (none or <3 months, 3 months to 3 years, 3 years to 5 years, more than 5 years), tubal ligation (yes/no) and parity (no children, 1 child, 2 children, 3 children, 4+ children).

Metabolites analyzed in this study belong to four metabolite groups (PC, LPC, SM and CER) previously hypothesized to be associated with ovarian cancer. Individual metabolites, the metabolite sums within metabolite groups, and the two created metabolite ratios showed a correlation structure corresponding to 4 groups. A Bonferroni adjusted p-value threshold of 0.0125 (0.05/4; 4 measured lipid groups) was used to evaluate statistical significance and to account for the effect of the four metabolite groups and their correlation structure on the results. We report statistically significant results (p-value≤0.0125) and nominally significant results (p-value≤0.05).

## Results

### Study population

Of the 252 cases analyzed, 176 cases were diagnosed with serous/PD tumors while 34 were classified as endometrioid/CC tumors (Table 1). The remaining cases were of mucinous or other types. Mean follow-up time was 12.3 years. Distributions of ovarian cancer risk factors were generally in the expected directions for cases and controls.

**Table 1:**
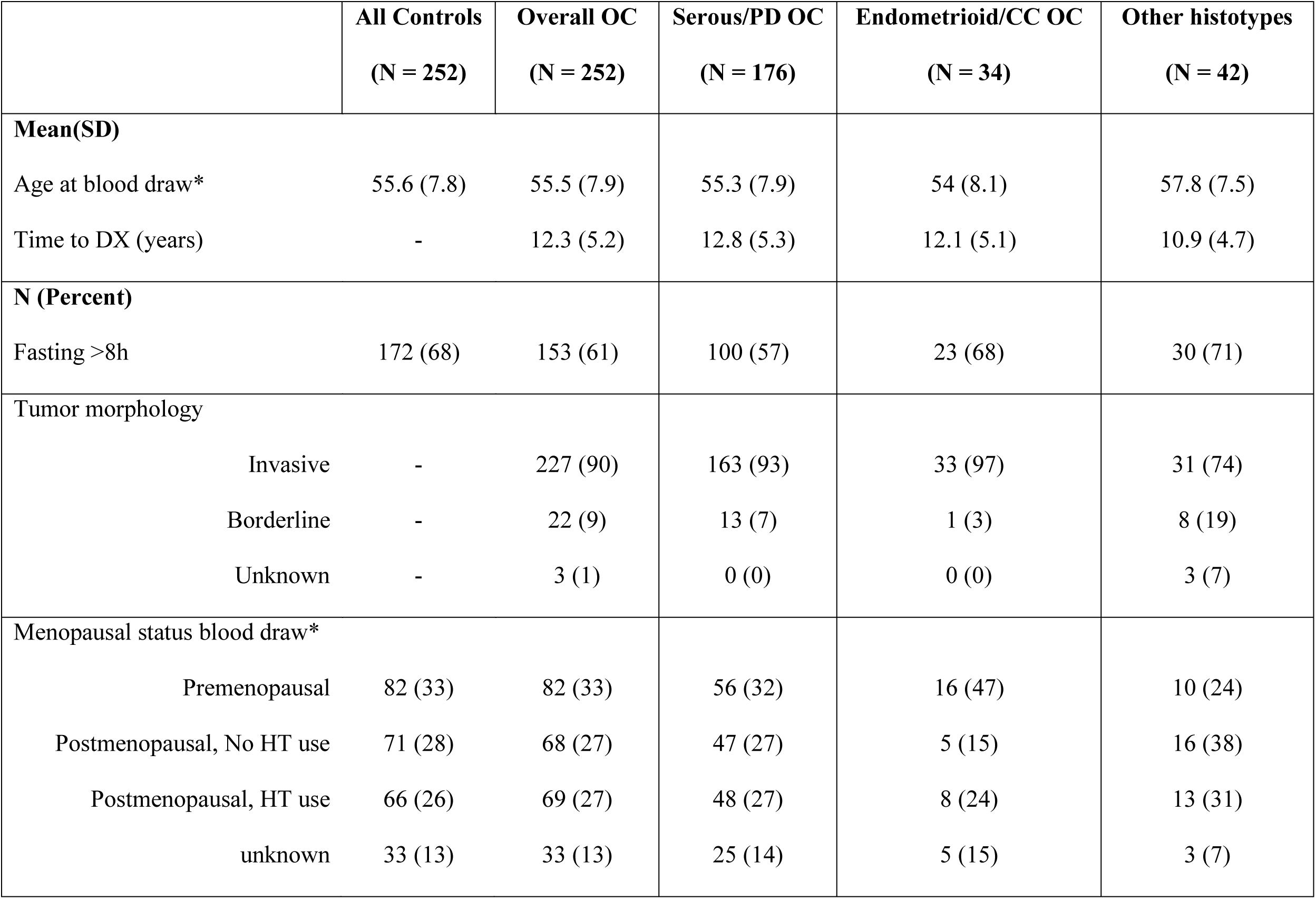
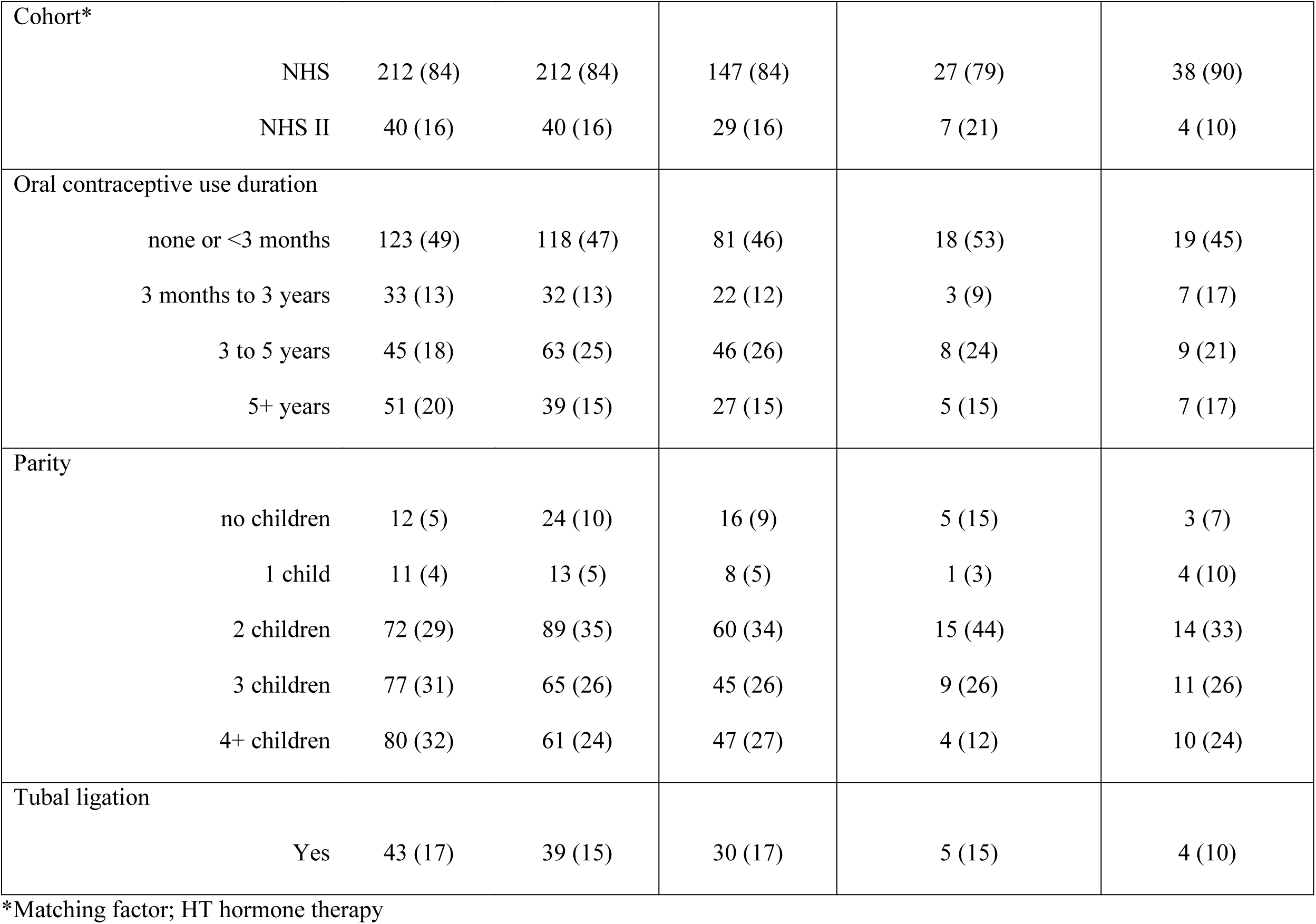
Characteristics of overall, serous/poorly differentiated (PD) and endometrioid/clear cell (CC) ovarian cancer (OC) cases, and all controls at time of blood collection.

|  | **All Controls**<br>**(N = 252)** | **Overall OC**<br>**(N = 252)** | **Serous/PD OC**<br>**(N = 176)** | **Endometrioid/CC OC**<br>**(N = 34)** | **Other histotypes**<br>**(N = 42)** |
| --- | --- | --- | --- | --- | --- |
| **Mean(SD)** |  |  |  |  |  |
| Age at blood draw* | 55.6 (7.8) | 55.5 (7.9) | 55.3 (7.9) | 54 (8.1) | 57.8 (7.5) |
| Time to DX (years) | - | 12.3 (5.2) | 12.8 (5.3) | 12.1 (5.1) | 10.9 (4.7) |
| **N (Percent)** |  |  |  |  |  |
| Fasting >8h | 172 (68) | 153 (61) | 100 (57) | 23 (68) | 30 (71) |
| Tumor morphology |  |  |  |  |  |
| Invasive | - | 227 (90) | 163 (93) | 33 (97) | 31 (74) |
| Borderline | - | 22 (9) | 13 (7) | 1 (3) | 8 (19) |
| Unknown | - | 3 (1) | 0 (0) | 0 (0) | 3 (7) |
| Menopausal status blood draw* |  |  |  |  |  |
| Premenopausal | 82 (33) | 82 (33) | 56 (32) | 16 (47) | 10 (24) |
| Postmenopausal, No HT use | 71 (28) | 68 (27) | 47 (27) | 5 (15) | 16 (38) |
| Postmenopausal, HT use | 66 (26) | 69 (27) | 48 (27) | 8 (24) | 13 (31) |
| unknown | 33 (13) | 33 (13) | 25 (14) | 5 (15) | 3 (7) |

|  |  |  |  |  |  |  |
| --- | --- | --- | --- | --- | --- | --- |
| Cohort* |  |  |  |  |  |  |
|  | NHS | 212 (84) | 212 (84) | 147 (84) | 27 (79) | 38 (90) |
|  | NHS II | 40 (16) | 40 (16) | 29 (16) | 7 (21) | 4 (10) |
| Oral contraceptive use duration |  |  |  |  |  |  |
|  | none or <3 months | 123 (49) | 118 (47) | 81 (46) | 18 (53) | 19 (45) |
|  | 3 months to 3 years | 33 (13) | 32 (13) | 22 (12) | 3 (9) | 7 (17) |
|  | 3 to 5 years | 45 (18) | 63 (25) | 46 (26) | 8 (24) | 9 (21) |
|  | 5+ years | 51 (20) | 39 (15) | 27 (15) | 5 (15) | 7 (17) |
| Parity |  |  |  |  |  |  |
|  | no children | 12 (5) | 24 (10) | 16 (9) | 5 (15) | 3 (7) |
|  | 1 child | 11 (4) | 13 (5) | 8 (5) | 1 (3) | 4 (10) |
|  | 2 children | 72 (29) | 89 (35) | 60 (34) | 15 (44) | 14 (33) |
|  | 3 children | 77 (31) | 65 (26) | 45 (26) | 9 (26) | 11 (26) |
|  | 4+ children | 80 (32) | 61 (24) | 47 (27) | 4 (12) | 10 (24) |
| Tubal ligation |  |  |  |  |  |  |
|  | Yes | 43 (17) | 39 (15) | 30 (17) | 5 (15) | 4 (10) |

\*Matching factor; HT hormone therapy

### Correlations between metabolites

Metabolites, metabolite sums, and metabolite ratios belonging to the same class were highly correlated with each other among controls (Figure 1), although the correlation among the SMs was slightly weaker than for the other classes. We observed somewhat lower correlations in postmenopausal women compared to premenopausal at blood draw (Supplementary Figure 1 and 2).

**Figure 1.**
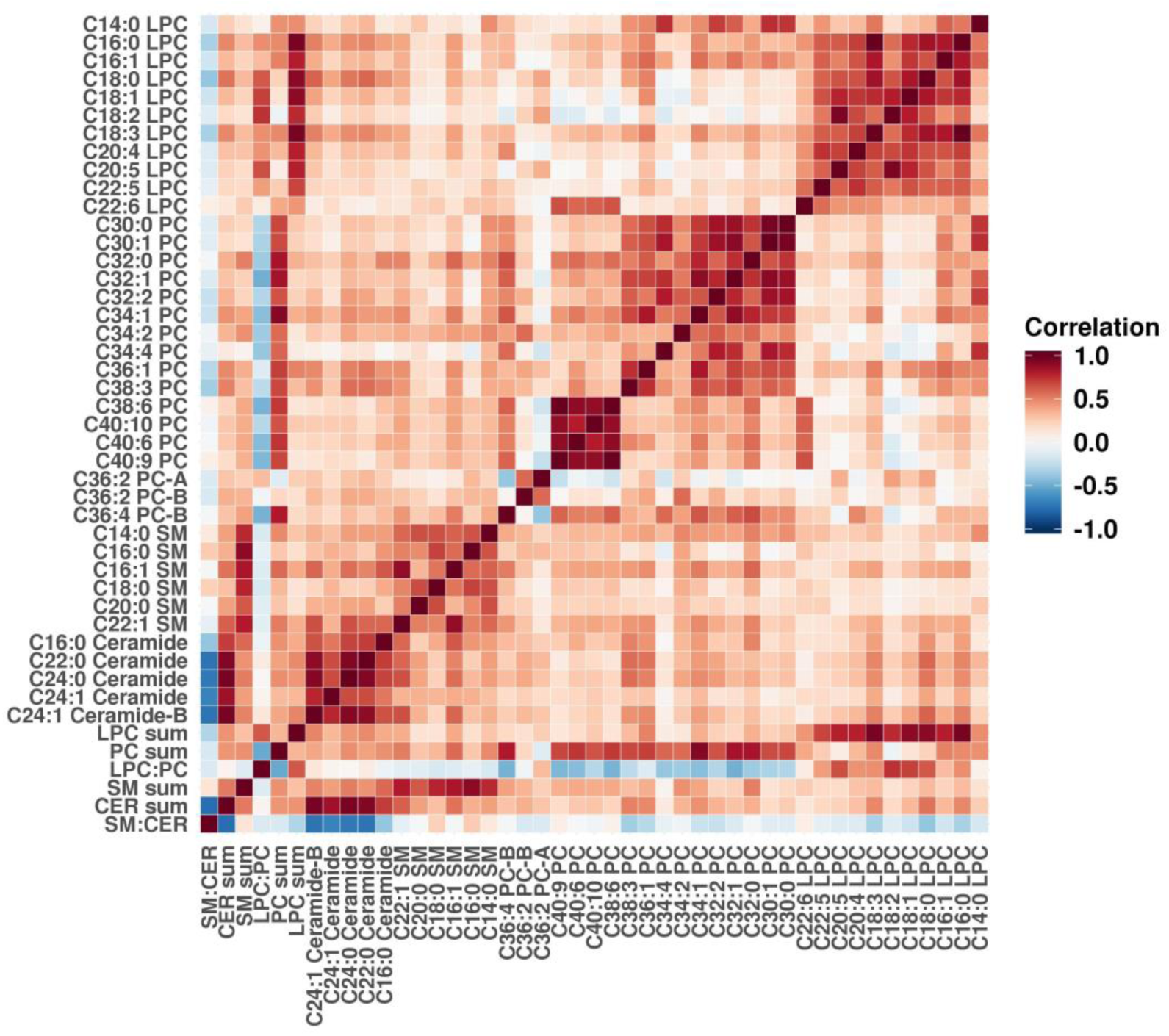
Metabolite correlations among controls. Pearson correlation was calculated for all pairs of individual metabolites, metabolite sums and metabolite ratios. Positive correlation coefficients are shown in shades of red and negative coefficients are shown in shades of blue.

### Metabolites associated with risk of ovarian cancer, serous/PD and endometrioid/CC ovarian cancer

Two individual SMs, C18:0 SM (OR (95%CI)=2.10 (1.26-3.49)) and C16:0 SM (OR=2.06), SM sum (OR=1.97), and C16:0 Ceramide (OR=1.95) were significantly positively associated with risk of overall ovarian cancer (Figure 2; Supplementary Table 1). No PCs or LPCs were significantly associated with risk of overall ovarian cancer. No metabolites were significantly associated with risk of serous/PD disease. However, C20:0 SM, C16:0 SM and the sum of all SMs were positively associated with risk of serous/PD ovarian cancer on the nominal scale. For endometrioid/CC disease, C18:1 LPC (OR (95%CI)=0.24 (0.07-0.69)) and LPC:PC (OR=0.24) were significantly inversely associated while C18:0 SM (OR=4.06) was significantly positively associated with risk.

**Figure 2.**
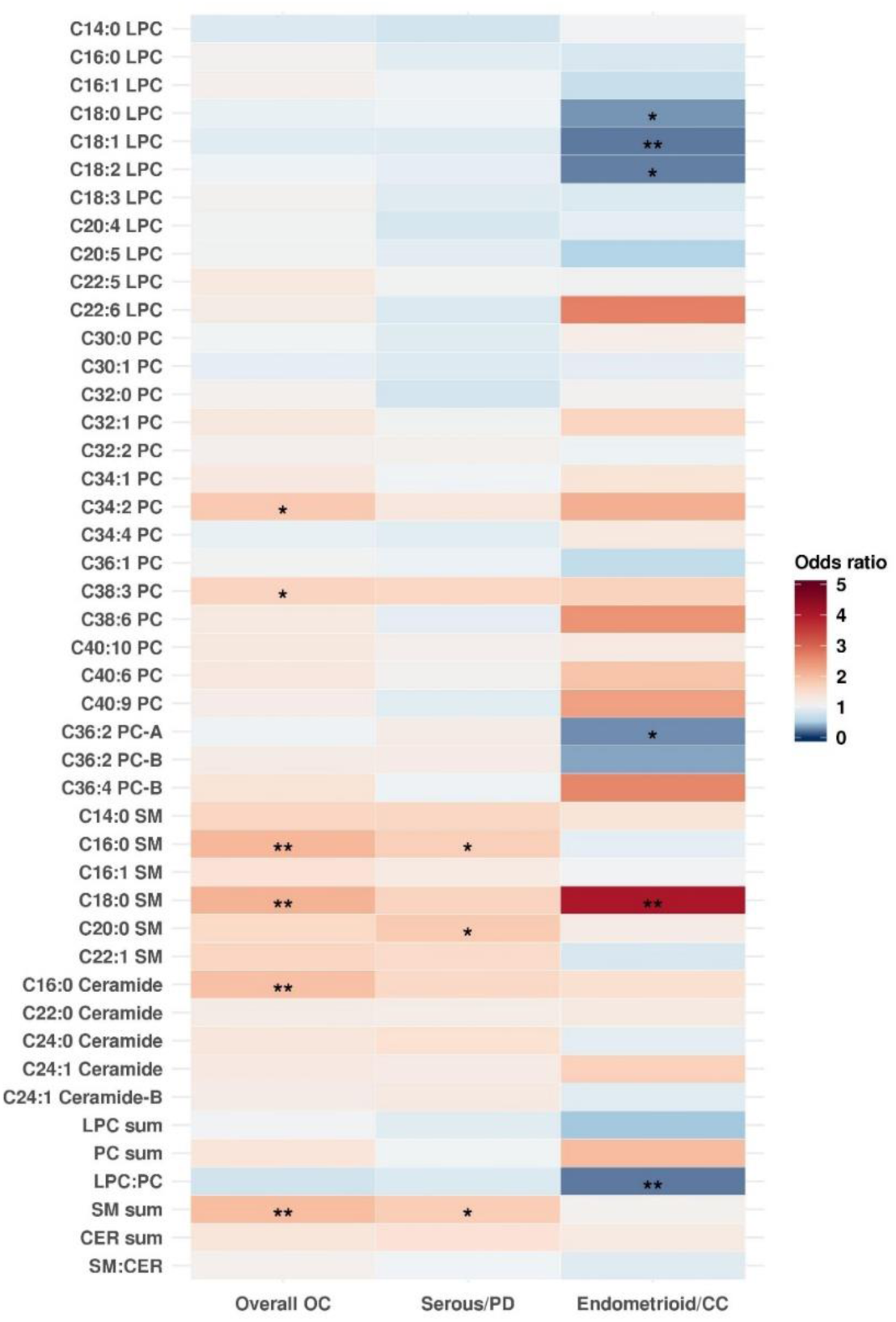
Odds ratios (OR) of risk of overall, serous/poorly differentiated and endometrioid/clear cell ovarian cancer for an increase from the 10^th^ to 90^th^ percentile in metabolite levels. OR>1 are shown in shades of red and OR<1 are shown in shades of blue. * p-value ≤0.05 (nominal significance), ** p-value ≤0.0125 (Bonferroni significance threshold). Estimates were adjusted for risk factors (duration of oral contraceptive use [none or <3 months, 3 months to 3 years, 3 years to 5 years, more than 5 years], tubal ligation [yes/no] and parity [no children, 1 child, 2 children, 3 children, 4+ children]) and additionally for matching factors (cohort [NHS, NHSII]; menopausal status and hormone therapy use at blood draw [premenopausal, postmenopausal and hormone therapy use, postmenopausal and no hormone therapy use, missing/unknown]; menopausal status at diagnosis [premenopausal, postmenopausal, or unknown]; age [±1 year], date of blood collection [± 1 month]; time of day of blood draw [±2 hours]; fasting status [>8 hours or ≤8 hours]) in subtype analyses.

### Metabolites associated with risk of rapidly fatal disease

C18:0 SM was significantly positively associated with risk of rapidly fatal disease OR (95% CI)=1.91 (1.19-3.09)) and less aggressive disease OR (95% CI)=2.10 (1.20-3.75) (Figure 3; Supplementary Table 2). C16:0 SM (OR=1.74), C16:0 CER (OR=1.77) and SM sum (OR=1.83) were nominally associated with rapidly fatal disease. C16:0 SM (OR=2.06) and SM sum (OR=1.88) were also nominally associated with less aggressive disease.

**Table 2:** Odds ratio (OR) of ovarian cancer, according to histotype, rapidly fatal status (death within 3 years of diagnosis versus not), menopausal status at blood draw, and time between blood collection and diagnosis, by quartiles (based on the distribution in controls) of the sum of all sphingomyelins.

|  |  | Quartile 1 | Quartile 2 | Quartile 3 | Quartile 4 | p-trend |
| --- | --- | --- | --- | --- | --- | --- |
|  |  | ( -2.88 -0.74 ] | ( -0.74 -0.12 ] | ( -0.12 0.54 ] | ( 0.54 2.88 ] |  |
| **Overall OC** | **OR (95%CI)** | REF | 0.95 (0.53,1.7) | 1.45 (0.83,2.51) | 1.67 (0.96,2.92) | 0.11 |
|  | **N controls** | 63 | 63 | 63 | 63 |  |
|  | **N cases** | 54 | 44 | 68 | 84 |  |
| **Serous/PD** | **OR (95%CI)** | REF | 1.02 (0.54,1.91) | 1.4 (0.77,2.57) | 1.58 (0.87,2.89) | 0.05 |
|  | **N controls** | 63 | 63 | 63 | 63 |  |
|  | **N cases** | 38 | 32 | 48 | 56 |  |
| **Endo/CC** | **OR (95%CI)** | REF | 1.35 (0.43,4.26) | 1.76 (0.61,5.29) | 0.88 (0.25,2.98) | 0.83 |
|  | **N controls** | 63 | 63 | 63 | 63 |  |
|  | **N cases** | 8 | 8 | 12 | 6 |  |
| **Rapidly fatal** | **OR (95%CI)** | REF | 0.91 (0.52,1.59) | 1.3 (0.76,2.23) | 1.63 (0.96,2.8) | 0.02 |
|  | **N controls** | 63 | 63 | 63 | 63 |  |
|  | **N cases** | 54 | 44 | 68 | 84 |  |
| **Less aggressive** | **OR (95%CI)** | REF | 0.93 (0.46,1.88) | 1.45 (0.76,2.8) | 1.83 (0.96,3.52) | 0.04 |
|  | **N controls** | 63 | 63 | 63 | 63 |  |
|  | **N cases** | 32 | 21 | 37 | 46 |  |
| **Premenopausal** | **OR (95%CI)** | REF | 0.68 (0.24,1.95) | 1.05 (0.38,2.87) | 0.42 (0.14,1.24) | 0.35 |
|  | **N controls** | 21 | 20 | 20 | 21 |  |
|  | **N cases** | 23 | 20 | 24 | 14 |  |
| **Postmenopausal** | **OR (95%CI)** | REF | 1.68 (0.77,3.64) | 1.89 (0.87,4.12) | 2.92 (1.36,6.27) | <0.01 |
|  | **N controls** | 35 | 34 | 34 | 34 |  |
|  | **N cases** | 21 | 28 | 33 | 54 |  |
| **Diagnosed 3-12**<br>**years after**<br>**blood draw** | **OR (95%CI)** | REF | 0.69 (0.32,1.46) | 1.36 (0.69,2.68) | 1.7 (0.88,3.36) | 0.04 |
|  | **N controls** | 63 | 63 | 63 | 63 |  |
|  | **N cases** | 29 | 17 | 34 | 40 |  |
| **Diagnosed 13-**<br>**23**<br>**years after**<br>**blood draw** | **OR (95%CI)** | REF | 1.19 (0.6,2.4) | 1.37 (0.7,2.72) | 1.75 (0.91,3.41) | 0.07 |
|  | **N controls** | 63 | 63 | 63 | 63 |  |
|  | **N cases** | 24 | 27 | 34 | 44 |  |

**Figure 3.**
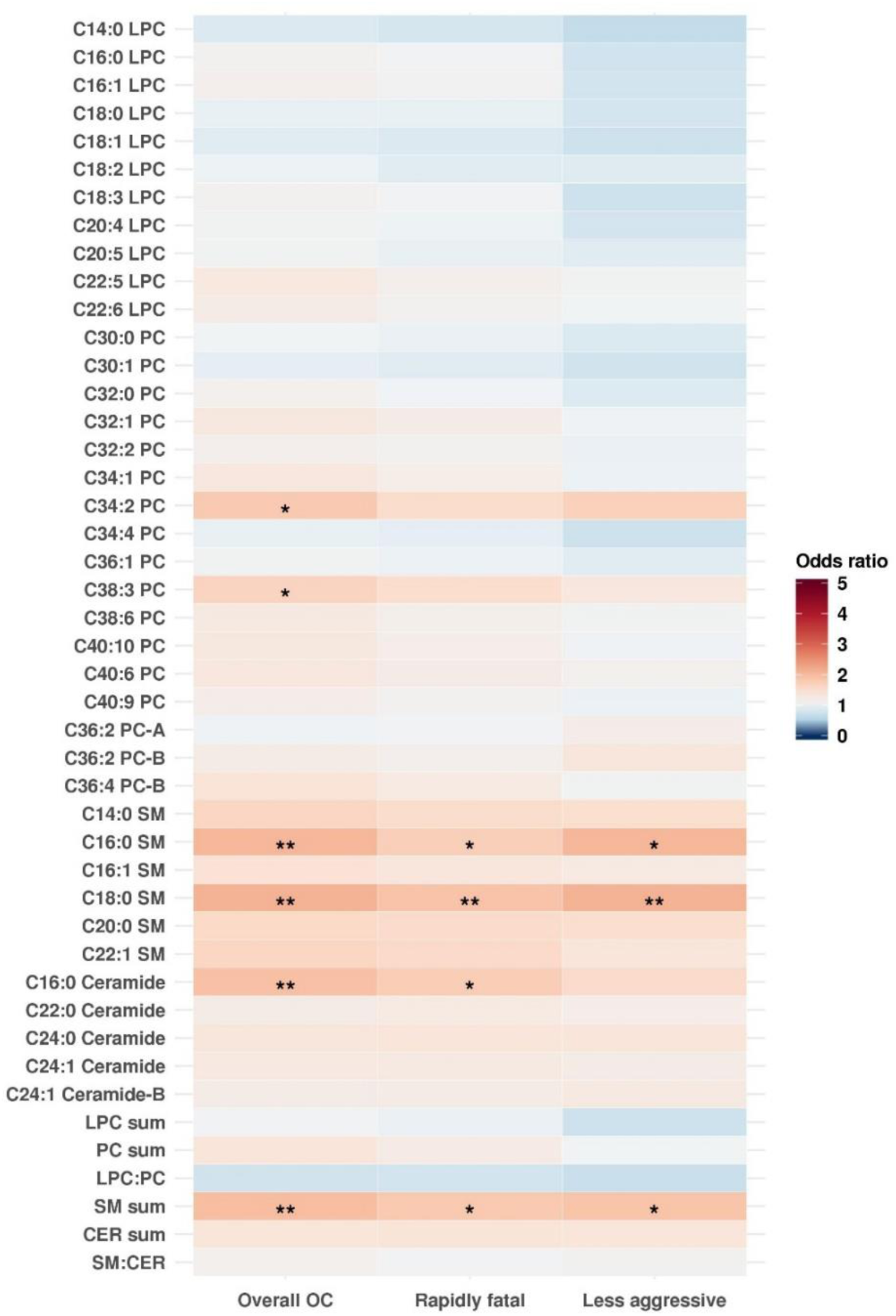
Odds ratios (OR) of risk of overall ovarian cancer and by rapidly fatal status for an increase from the 10^th^ to 90^th^ percentile in metabolite levels. OR>1 are shown in shades of red and OR<1 are shown in shades of blue. * p-value ≤0.05 (nominal significance), ** p-value ≤0.0125 (Bonferroni significance threshold). Estimates were adjusted for risk factors (duration of oral contraceptive use [none or <3 months, 3 months to 3 years, 3 years to 5 years, more than 5 years], tubal ligation [yes/no] and parity [no children, 1 child, 2 children, 3 children, 4+ children]) and additionally for matching factors (cohort [NHS, NHSII]; menopausal status and hormone therapy use at blood draw [premenopausal, postmenopausal and hormone therapy use, postmenopausal and no hormone therapy use, missing/unknown]; menopausal status at diagnosis [premenopausal, postmenopausal, or unknown]; age [±1 year], date of blood collection [± 1 month]; time of day of blood draw [±2 hours]; fasting status [>8 hours or ≤8 hours]) in subtype analyses.

### Metabolites associated with risk of ovarian cancer by menopausal status at blood draw

Multiple markers were significantly positively associated with risk of ovarian cancer among postmenopausal women at blood collection (Figure 4; Supplementary Table 3): SM sum (OR (95%CI)=3.22 (1.51-6.86)), C22:1 SM (OR=3.11), and C16:0 SM (OR= 2.98). Of the six measured SMs, five were positively associated with risk of ovarian cancer (2 significantly and 3 nominally), while SM sum was significantly positively associated with risk among postmenopausal women at blood collection. No metabolites were significantly associated with risk of overall ovarian cancer among premenopausal women at blood collection. However, 5 out of 6 SMs and SM sum had ORs below 1.0, although not significant even on the nominal scale.

**Figure 4.**
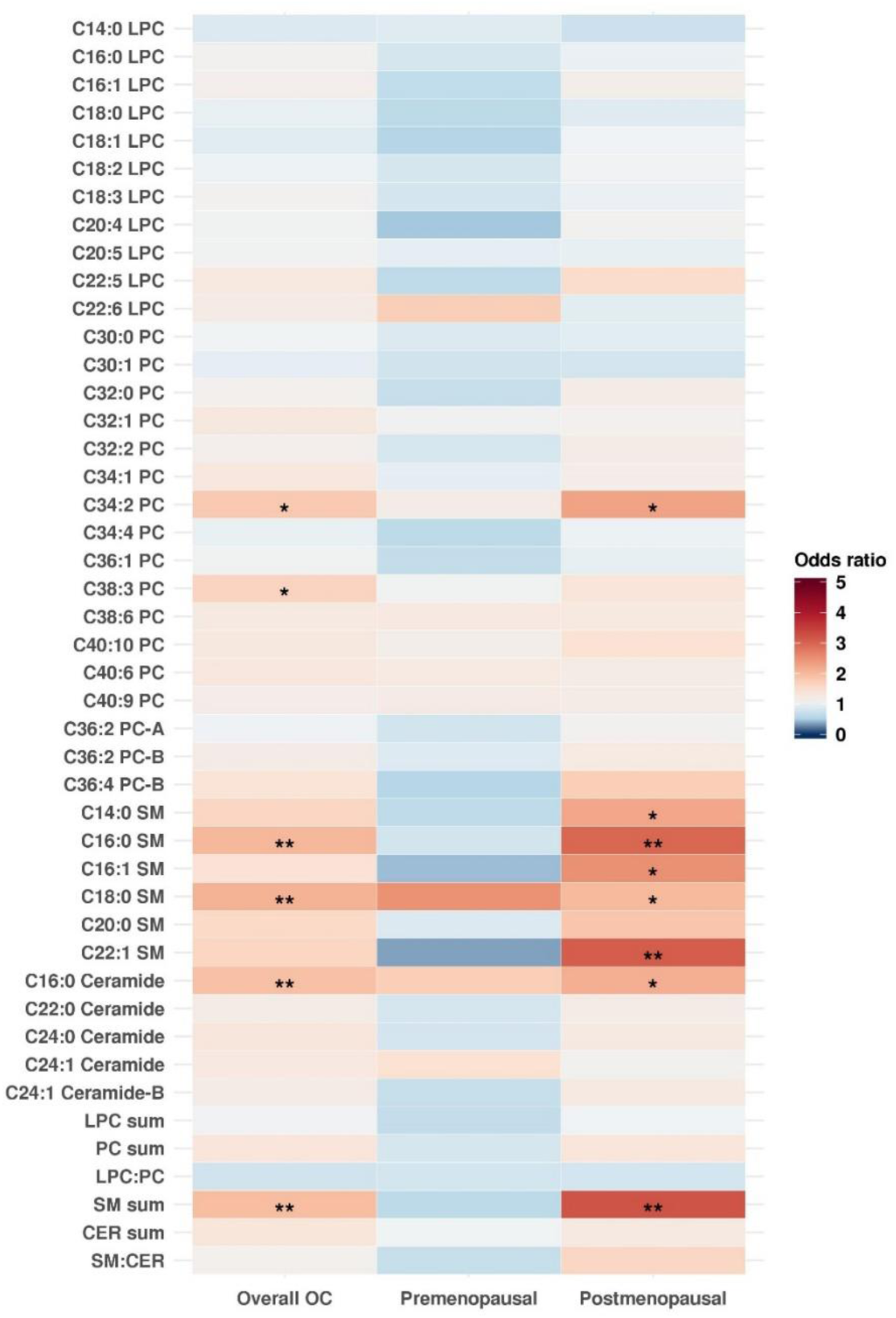
Odds ratios (OR) of risk of overall ovarian cancer and by menopausal status at blood draw for an increase from the 10^th^ to 90^th^ percentile in metabolite levels. OR>1 are shown in shades of red and OR<1 are shown in shades of blue. * p-value ≤0.05 (nominal significance), ** p-value ≤0.0125 (Bonferroni significance threshold). Estimates were adjusted for risk factors (duration of oral contraceptive use [none or <3 months, 3 months to 3 years, 3 years to 5 years, more than 5 years], tubal ligation [yes/no] and parity [no children, 1 child, 2 children, 3 children, 4+ children]) and additionally for matching factors (cohort [NHS, NHSII]; menopausal status and hormone therapy use at blood draw [premenopausal, postmenopausal and hormone therapy use, postmenopausal and no hormone therapy use, missing/unknown]; menopausal status at diagnosis [premenopausal, postmenopausal, or unknown]; age [±1 year], date of blood collection [± 1 month]; time of day of blood draw [±2 hours]; fasting status [>8 hours or ≤8 hours]) in subtype analyses.

### Metabolites associated with risk of ovarian cancer by time elapsed between blood draw to diagnosis

In general, risk estimates for the individual SMs and their sum were in the same direction (positive) for cases diagnosed 3-12 years after blood draw versus 13-23 years after blood draw (Supplementary Figure 3; Supplementary Table 4). Specifically, C20:0 SM (OR (95%CI)=2.18 (1.22-3.94)) was significantly positively associated with risk of ovarian cancer thirteen to twenty-three years after blood draw, and C18:0 SM was nominally significant. C34:2 PC (OR=2.24) was significantly positively associated with risk of ovarian cancer three to twelve years after blood draw. Additional six metabolites (1 PC, 2SMs, 1 CER, PC sum and SM sum) were nominally positively associated with ovarian cancer risk in this period. Notably, the risk estimates for the 2 PCs and PC sum were positively associated with ovarian cancer developing in 3-12 years after blood draw, but the risk estimates for these factors were null or non-significantly inverse for cancer developing 13-23 years after blood draw.

### Sphingomyelins

To better understand the dose-response relationship between SM (analyzed as the sum of all measured SMs) and risk of ovarian cancer, we calculated odds ratios for the three highest quartiles compared to the first quartile (Table 2). We observed a dose-response association for overall ovarian cancer among postmenopausal women at blood draw with OR of 1.68 (95%CI=0.77-3.64) for quartile 2, 1.89 (95%CI=0.87-4.12) for quartile 3, and 2.92 (95%CI=1.36-6.27) to the lowest quartile, with a p-trend <0.01. Nominally significant trends (OR, top vs. bottom quartile) were observed for overall ovarian cancer (OR=1.67), serous/PD disease (OR=1.58), both rapidly fatal (OR=1.63) and less aggressive tumors (OR=1.83). Similar ORs comparing the top vs. bottom quartile of total SMs were observed for cases diagnosed 3-12 years (OR=1.70; p-trend=0.04) and 13-23 years after blood draw (OR=1.75, p-trend=0.07).

## Discussion

This is the first large-scale prospective study to examine the relationship of circulating LPCs, PCs, SMs and ceramides with risk of ovarian cancer. Higher levels of two circulating SMs and a ceramide were significantly associated with about a 2-fold increased risk of ovarian cancer when metabolites levels increased from 10^th^ to 90^th^ percentile of the distribution. In addition to individual SMs, the sum of all measured SMs was also associated with a 2-fold increased risk of ovarian cancer. While no metabolites met the Bonferroni threshold for significance for risk of serous/PD tumors, the SM sum and 2 SM were nominally significant. Strength of associations tended to be stronger for endometrioid/CC tumors: one SM was significantly associated with a 4.44-fold increased while one LPC and LPC:PC were significantly associated with 0.2-fold lower risk of endometrioid/CC tumors for an increase in levels from the 10^th^ to 90^th^ percentile, although the overall sample size was limited for this subtype. SMs and SM sum showed similar ORs when comparing rapidly fatal with less aggressive tumors. Notably, more associations were observed among women who were postmenopausal at blood draw, with a nearly 3-fold increased risk for the top versus bottom quartile of the SM sum, representing a potentially novel and strong risk factor.

The only other prospective study investigating these four groups of lipids, assessed their associations with risk of three cancers (40). Similar to our results for ovarian cancer risk, positive associations, although not significant, were observed for several SMs with risk of breast cancer (C16:0 SM, C18:0 SM, C20:2 SM, and C24:0; 4/7 measured SM) (40). Additionally, two of seven measured SMs were nominally positively associated with risk of prostate cancer and five of seven SMs were nominally positively associated with colorectal cancer.

SMs are the most abundant class of sphingolipids in the cell, essential elements of the cell membrane, and critical players in cell function. SM are *de novo* synthesized from ceramides and hydrolyzed back into ceramides, which are involved in cellular proliferation, growth and apoptosis (41). SMs were higher in ovarian tumors compared to healthy tissue (33, 42). Increased levels of SMs were also reported in taxol-resistant human ovarian cancer cell lines (43). Additionally, acid sphingomyelinase (enzyme converting SMs into ceramides) has been associated with cisplatin-resistance in ovarian tumors (44) and was lower in human ovarian cancer cells compared to human primary ovarian cells (45). A recent study showed that sphingomyelin in microdomains of the plasma membrane regulates amino acid-stimulated mTOR signal activation (46), which is essential for cell growth and proliferation (47). Adding to the already existing literature, we show that circulating levels of SMs are higher in future ovarian cancer cases compared to healthy controls up to 23 years before diagnosis.

Interestingly, Kuehn and colleagues (40) found C18:0 LPC to be associated with lower risk of breast, prostate, and colon cancers (40). C18:0 LPC was inversely associated with risk of endometrioid/CC tumors in our analysis. Furthermore, we observed inverse associations for C18:1 LPC and C18:2 LPC with risk of endometrioid/CC tumors, with C18:1 LPC and C18:2 LPC being strongly correlated with C18:0 LPC. A risk factor for endometrioid/CC tumors is a diagnosis of endometriosis. In a recent lipidomic study of endometrial fluid in women with ovarian endometriosis, lysophospholipids, calculated as the sum of LPCs and LPEs, were found to be lower in cases compared to healthy controls (48). As LPEs were not part of our analyses, it is difficult to say whether these two results are in agreement.

Additionally, the lipidomics study found ceramides to be down-regulated in endometriosis cases. In our analysis, two of the five measured ceramides were associated with lower risk of endometrioid/CC and three ceramides were associated with lower risk of ovarian cancer in premenopausal women, who were younger at blood collection, and represent the most similar subgroup to the women in the endometriosis study. Larger studies are needed to assess ceramides and LPCs as potential risk biomarkers for endometrioid tumors, particularly in younger women.

Additionally, we observed several similar patterns of associations by lipid group, although not significant, comparing our results with the results from Kuehn et al. (40). In addition to SMs, LPCs were nominally associated with lower risk of breast, colorectal, prostate, endometrioid/CC ovarian tumors and serous/PD ovarian tumors. PCs were nominally associated with higher risk of colorectal, breast and endometrioid/PD ovarian tumors. These findings suggest that systematic lipid changes observed in several cancer cells and tumors (49) may be measured in blood samples 3-23 years before diagnosis.

Our study has several strengths and limitations. To the best of our knowledge, this is the first study assessing the associations of these lipids with ovarian cancer risk. Even though our power was limited we conducted exploratory analyses by histotype, menopausal status, rapidly fatal status and time between blood draw and diagnosis. Given the low incidence rate of ovarian cancer, our study has a relatively large sample size with 252 cases and 252 matched controls. Additional strengths include the long follow-up time and detailed covariate information. Another limitation is represented by the one-point-in-time blood samples analyzed here. To address this limitation, we conducted a pilot study showing that the majority of the measured metabolites have a high within person stability over time (ICC or Spearman correlations were higher than 0.7 for CER, 0.68 for LPC, 0.65 for PC and 0.76 for SM) (39).

In summary, we found SMs to be associated with increased risk of ovarian cancer, particularly among postmenopausal women. We observed elevated levels of SMs in ovarian cancer cases 3-23 years before diagnosis. Additionally, C18:1 LPC and LPC:PC was associated with lower risk of endometrioid/CC tumors. These results provide new and promising candidates for risk and early detection biomarkers. Experimental studies may help identify the mechanisms through which a dysregulated lipid metabolism supports carcinogenesis, potentially leading to the development of new therapeutic targets for prevention and treatment of ovarian cancer. Population studies are needed to validate these findings and further characterize their potential as risk and early detection biomarkers. If confirmed, these markers may be used to identify high-risk women who can benefit from preventive interventions. Additional studies are needed to identify biological predictors of circulating SMs, the underlying biological relationship with ovarian tumors and the ovaries and ultimately how high doses promote carcinogenesis.

## Acknowledgements

We would like to thank the participants and staff of the NHS and NHS II for their valuable contributions as well as the following state cancer registries for their help: AL, AZ, AR, CA, CO, CT, DE, FL, GA, ID, IL, IN, IA, KY, LA, ME, MD, MA, MI, NE, NH, NJ, NY, NC, ND, OH, OK, OR, PA, RI, SC, TN, TX, VA, WA, WY. The authors assume full responsibility for analyses and interpretation of these data.

## Notes

The authors have no conflicts of interest to report.

## References

1. American Cancer Society. Cancer Facts & Figures 2018. Atlanta: American Cancer Society; 2018.

2. Röhrig F, Schulze A. The multifaceted roles of fatty acid synthesis in cancer. Nat Rev Cancer. 2016;16(11):732.

3. Pyragius CE, Fuller M, Ricciardelli C, Oehler MK. Aberrant lipid metabolism: an emerging diagnostic and therapeutic target in ovarian cancer. Int J Mol Sci. 2013;14(4):7742–56.

4. Tania M, Khan M, Song Y. Association of lipid metabolism with ovarian cancer. Current oncology. 2010;17(5):6.

5. Helzlsouer KJ, Alberg AJ, Norkus EP, Morris JS, Hoffman SC, Comstock GW. Prospective study of serum micronutrients and ovarian cancer. J Natl Cancer Inst. 1996;88(1):32–7.

6. Melvin JC, Seth D, Holmberg L, Garmo H, Hammar N, Jungner I, et al. Lipid profiles and risk of breast and ovarian cancer in the Swedish AMORIS study. Cancer Epidemiol Biomarkers Prev. 2012;21(8):1381–4.

7. Braicu EI, Darb-Esfahani S, Schmitt WD, Koistinen KM, Heiskanen L, Pöhö P, et al. High-grade ovarian serous carcinoma patients exhibit profound alterations in lipid metabolism. Oncotarget. 2017;8(61):102912.

8. Ke C, Hou Y, Zhang H, Fan L, Ge T, Guo B, et al. Large-scale profiling of metabolic dysregulation in ovarian cancer. Int J Cancer. 2015;136(3):516–26.

9. Ke C, Li A, Hou Y, Sun M, Yang K, Cheng J, et al. Metabolic phenotyping for monitoring ovarian cancer patients. Sci Rep. 2016;6:23334.

10. Xu Y, Fang X, Casey G, Mills G. Lysophospholipids activate ovarian and breast cancer cells. Biochem J. 1995;309(3):933–40.

11. Fang X, Gaudette D, Furui T, Mao M, Estrella V, Eder A, et al. Lysophospholipid growth factors in the initiation, progression, metastases, and management of ovarian cancer. Ann N Y Acad Sci. 2000;905(1):188–208.

12. Fang X, Yu S, Bast RC, Liu S, Xu H-J, Hu S-X, et al. Mechanisms for lysophosphatidic acid-induced cytokine production in ovarian cancer cells. J Biol Chem. 2004;279(10):9653–61.

13. Sawada K, Morishige K-i, Tahara M, Kawagishi R, Ikebuchi Y, Tasaka K, et al. Alendronate inhibits lysophosphatidic acid-induced migration of human ovarian cancer cells by attenuating the activation of rho. Cancer Res. 2002;62(21):6015–20.

14. Cai Q, Zhao Z, Antalis C, Yan L, Del Priore G, Hamed AH, et al. Elevated and secreted phospholipase A2 activities as new potential therapeutic targets in human epithelial ovarian cancer. The FASEB Journal. 2012;26(8):3306–20.

15. Tokumura A, Kume T, Fukuzawa K, Tahara M, Tasaka K, Aoki J, et al. Peritoneal fluids from patients with certain gynecologic tumor contain elevated levels of bioactive lysophospholipase D activity. Life Sci. 2007;80(18):1641–9.

16. Pyragius CE, Fuller M, Ricciardelli C, Oehler MK. Aberrant lipid metabolism: an emerging diagnostic and therapeutic target in ovarian cancer. Int J Mol Sci. 2013;14(4):7742–56.

17. Cai Q, Zhao Z, Antalis C, Yan L, Del Priore G, Hamed AH, et al. Elevated and secreted phospholipase A(2) activities as new potential therapeutic targets in human epithelial ovarian cancer. Faseb J. 2012;26(8):3306–20.

18. Fan L, Zhang W, Yin M, Zhang T, Wu X, Zhang H, et al. Identification of metabolic biomarkers to diagnose epithelial ovarian cancer using a UPLC/QTOF/MS platform. Acta Oncol. 2012;51(4):473–9.

19. Okita M, Gaudette DC, Mills GB, Holub BJ. Elevated levels and altered fatty acid composition of plasma lysophosphatidylcholine(lysoPC) in ovarian cancer patients. Int J Cancer. 1997;71(1):31–4.

20. Sedlakova I, Vavrova J, Tosner J, Hanousek L. Lysophosphatidic acid: an ovarian cancer marker. Eur J Gynaecol Oncol. 2008;29(5):511–4.

21. Sedlakova I, Vavrova J, Tosner J, Hanousek L. Lysophosphatidic acid (LPA)-a perspective marker in ovarian cancer. Tumour Biol. 2011;32(2):311–6.

22. Shan L, Chen YA, Davis L, Han G, Zhu W, Molina AD, et al. Measurement of phospholipids may improve diagnostic accuracy in ovarian cancer. PLoS One. 2012;7(10):e46846.

23. Sutphen R, Xu Y, Wilbanks GD, Fiorica J, Grendys EC, Jr., LaPolla JP, et al. Lysophospholipids are potential biomarkers of ovarian cancer. Cancer Epidemiol Biomarkers Prev. 2004;13(7):1185–91.

24. Xiao YJ, Schwartz B, Washington M, Kennedy A, Webster K, Belinson J, et al. Electrospray ionization mass spectrometry analysis of lysophospholipids in human ascitic fluids: comparison of the lysophospholipid contents in malignant vs nonmalignant ascitic fluids. Anal Biochem. 2001;290(2):302–13.

25. Xu Y, Shen Z, Wiper DW, Wu M, Morton RE, Elson P, et al. Lysophosphatidic acid as a potential biomarker for ovarian and other gynecologic cancers. Jama. 1998;280(8):719–23.

26. Zhang T, Wu X, Ke C, Yin M, Li Z, Fan L, et al. Identification of potential biomarkers for ovarian cancer by urinary metabolomic profiling. J Proteome Res. 2013;12(1):505–12.

27. Lin HY, Delmas D, Vang O, Hsieh TC, Lin S, Cheng GY, et al. Mechanisms of ceramide-induced COX-2-dependent apoptosis in human ovarian cancer OVCAR-3 cells partially overlapped with resveratrol. J Cell Biochem. 2013;114(8):1940–54.

28. Morita Y, Perez GI, Paris F, Miranda SR, Ehleiter D, Haimovitz-Friedman A, et al. Oocyte apoptosis is suppressed by disruption of the acid sphingomyelinase gene or by sphingosine-1-phosphate therapy. Nat Med. 2000;6(10):1109–14.

29. Tilly JL, Kolesnick RN. Sphingolipids, apoptosis, cancer treatments and the ovary: investigating a crime against female fertility. Biochim Biophys Acta. 2002;1585(2-3):135–8.

30. Prinetti A, Millimaggi D, D’Ascenzo S, Clarkson M, Bettiga A, Chigorno V, et al. Lack of ceramide generation and altered sphingolipid composition are associated with drug resistance in human ovarian carcinoma cells. Biochem J. 2006;395(2):311–8.

31. Kitatani K, Usui T, Sriraman SK, Toyoshima M, Ishibashi M, Shigeta S, et al. Ceramide limits phosphatidylinositol-3-kinase C2β-controlled cell motility in ovarian cancer: potential of ceramide as a metastasis-suppressor lipid. Oncogene. 2016;35(21):2801.

32. Ogretmen B, Hannun YA. Biologically active sphingolipids in cancer pathogenesis and treatment. Nat Rev Cancer. 2004;4(8):604–16.

33. Fong MY, McDunn J, Kakar SS. Identification of metabolites in the normal ovary and their transformation in primary and metastatic ovarian cancer. PLoS One. 2011;6(5):e19963.

34. Hankinson SE, Willett WC, Michaud DS, Manson JE, Colditz GA, Longcope C, et al. Plasma prolactin levels and subsequent risk of breast cancer in postmenopausal women. J Natl Cancer Inst. 1999;91(7):629–34.

35. Tworoger SS, Sluss P, Hankinson SE. Association between plasma prolactin concentrations and risk of breast cancer among predominately premenopausal women. Cancer research. 2006;66(4):2476–82.

36. Mascanfroni ID, Takenaka MC, Yeste A, Patel B, Wu Y, Kenison JE, et al. Metabolic control of type 1 regulatory T cell differentiation by AHR and HIF1-α. Nat Med. 2015;21(6):638.

37. O’sullivan JF, Morningstar JE, Yang Q, Zheng B, Gao Y, Jeanfavre S, et al. Dimethylguanidino valeric acid is a marker of liver fat and predicts diabetes. The Journal of clinical investigation. 2017;127(12):4394–402.

38. Paynter NP, Balasubramanian R, Giulianini F, Wang DD, Tinker LF, Gopal S, et al. Metabolic predictors of incident coronary heart disease in women. Circulation. 2018;137(8):841–53.

39. Townsend MK, Clish CB, Kraft P, Wu C, Souza AL, Deik AA, et al. Reproducibility of metabolomic profiles among men and women in 2 large cohort studies. Clin Chem. 2013;59(11):1657–67.

40. Kühn T, Floegel A, Sookthai D, Johnson T, Rolle-Kampczyk U, Otto W, et al. Higher plasma levels of lysophosphatidylcholine 18: 0 are related to a lower risk of common cancers in a prospective metabolomics study. BMC Med. 2016;14(1):13.

41. Bienias K, Fiedorowicz A, Sadowska A, Prokopiuk S, Car H. Regulation of sphingomyelin metabolism. Pharmacol Rep. 2016;68(3):570–81.

42. Kozar N, Kruusmaa K, Bitenc M, Argamasilla R, Adsuar A, Goswami N, et al. Metabolomic profiling suggests long chain ceramides and sphingomyelins as a possible diagnostic biomarker of epithelial ovarian cancer. Clin Chim Acta. 2018;481:108–14.

43. Huang H, Tong T-T, Yau L-F, Chen C-Y, Mi J-N, Wang J-R, et al. LC-MS Based Sphingolipidomic Study on A2780 Human Ovarian Cancer Cell Line and its Taxol-resistant Strain. Sci Rep. 2016;6:34684.

44. Maurmann L, Belkacemi L, Adams N, Majmudar P, Moghaddas S, Bose R. A novel cisplatin mediated apoptosis pathway is associated with acid sphingomyelinase and FAS proapoptotic protein activation in ovarian cancer. Apoptosis. 2015;20(7):960–74.

45. Dai S, Liu J, Sun X, Wang N. Acid sphingomyelinase, a novel negative biomarker of ovarian cancer. Eur Rev Med Pharmacol Sci. 2015;19(11):2076–83.

46. Zama K, Mitsutake S, Okazaki T, Igarashi Y. Sphingomyelin in microdomains of the plasma membrane regulates amino acid-stimulated mTOR signal activation. Cell Biol Int. 2018.

47. Wullschleger S, Loewith R, Hall MN. TOR signaling in growth and metabolism. Cell. 2006;124(3):471–84.

48. Domínguez F, Ferrando M, Díaz-Gimeno P, Quintana F, Fernández G, Castells I, et al. Lipidomic profiling of endometrial fluid in women with ovarian endometriosis. Biol Reprod. 2017;96(4):772–9.

49. Ray U, Roy SS. Aberrant lipid metabolism in cancer cells–the role of oncolipid-activated signaling. The FEBS journal. 2018;285(3):432–43.

