## Supplementary Materials for "Circulating Lysophosphatidylcholines, Phosphatidylcholines, Ceramides, and Sphingomyelins and Ovarian Cancer Risk: a 23-year Prospective Study"

### Study Population

Blood collection: Women arranged to have their blood drawn and shipped with an ice pack, via overnight courier, to our laboratory, where it was processed and separated into plasma, red blood cell, and white blood cell components and frozen in gasketed cryovials in the vapor phase of liquid nitrogen freezers. Between 1996 and 1999, 29,611 NHSII participants provided blood samples and completed a short questionnaire . Premenopausal women (*n*=18,521) who had not taken hormones, been pregnant, or lactated within the past 6 months provided blood samples drawn 7–9 days before the anticipated start of their next menstrual cycle (luteal phase). Other women (*n* = 11,090) provided a single 30-mL untimed blood sample. Samples were shipped and processed identically to the NHS samples.

Incident cases of epithelial ovarian cancer were identified through the biennial questionnaires or via linkage with the National Death Index. For women reporting a new ovarian cancer diagnosis or cases identified through death certificates, we obtained related medical records and pathology reports; for cases who had died we linked to the relevant cancer registry is medical records were unattainable. A gynecologic pathologist reviewed the records to confirm the diagnosis and abstract date of diagnosis, invasiveness, stage, and histologic subtype (serous, poorly differentiated [PD], endometrioid, clear cell [CC], mucinous, other/unknown). Date of death was extracted from the death certificate. Participants diagnosed with invasive disease and who died within 3 years of diagnosis were defined as rapidly fatal cases.

### Metabolite Profiling

For each method, pooled plasma reference samples were included every 20 samples and results were standardized using the ratio of the value of the sample to the value of the nearest pooled reference multiplied by the median of all reference values for the metabolite. Samples from the two cohorts were run together, with matched case-control pairs distributed randomly within the batch, and the order of the case and controls within each pair was also randomly designated. Therefore, the case and its control were always directly adjacent to each other in the analytic run, thereby limiting variability in platform performance across matched case-control pairs. In addition to the participants' samples, 64 quality control (QC) samples, to which the laboratory was blinded, were randomly inserted in pairs among participant samples and profiled.

Hydrophilic interaction liquid chromatography (HILIC) analyses of water- soluble metabolites in the positive ionization mode (HILIC-positive) were conducted using an LC-MS system comprised of a Shimadzu Nexera X2 U-HPLC (Shimadzu Corp.; Marlborough, MA) coupled to a Q Exactive mass spectrometer (Thermo Fisher Scientific; Waltham, MA). Metabolites were extracted from plasma (10 µL) using 90 µL of acetonitrile/methanol/formic acid (74.9:24.9:0.2 v/v/v) containing stable isotope-labeled internal standards (valine-d8, Sigma-Aldrich; St. Louis, MO; and phenylalanine-d8, Cambridge Isotope Laboratories; Andover, MA). The samples were centrifuged (10 min, 9,000 x g, 4°C), and the supernatants were injected directly onto a 150 x 2 mm, 3 µm Atlantis HILIC column (Waters; Milford, MA). The column was eluted isocratically at a flow rate of 250 µL/min with 5% mobile phase A (10 mM ammonium formate and 0.1% formic acid in water) for 0.5 minute followed by a linear gradient to 40% mobile phase B (acetonitrile with 0.1% formic acid) over 10 minutes. MS analyses were carried out using electrospray ionization in the positive ion mode using full scan analysis over 70-800 m/z at 70,000 resolution and 3 Hz data acquisition rate. Other MS settings were: sheath gas 40, sweep gas 2, spray voltage 3.5 kV, capillary temperature 350°C, S-lens RF 40, heater temperature 300°C, microscans 1, automatic gain control target 1e6, and maximum ion time 250 ms.

Plasma lipids were profiled using a Shimadzu Nexera X2 U-HPLC (Shimadzu Corp.; Marlborough, MA) (C8-positive). Lipids were extracted from plasma (10 µL) using 190 µL of isopropanol containing 1,2-didodecanoyl-sn-glycero-3-phosphocholine (Avanti Polar Lipids; Alabaster, AL). After centrifugation, supernatants were injected directly onto a 100 x 2.1 mm, 1.7 µm ACQUITY BEH C8 column (Waters; Milford, MA). The column was eluted isocratically with 80% mobile phase A (95:5:0.1 vol/vol/vol 10mM ammonium acetate/methanol/formic acid) for 1 minute followed by a linear gradient to 80% mobile-phase B (99.9:0.1 vol/vol methanol/formic acid) over 2 minutes, a linear gradient to 100% mobile phase B over 7 minutes, then 3 minutes at 100% mobile-phase B. MS analyses were carried out using electrospray ionization in the positive ion mode using full scan analysis over 200–1100 m/z at 70,000 resolution and 3 Hz data acquisition rate. Other MS settings were: sheath gas 50, in source CID 5 eV, sweep gas 5, spray voltage 3 kV, capillary temperature 300°C, S-lens RF 60, heater temperature 300°C, microscans 1, automatic gain control target 1e6, and maximum ion time 100 ms. Lipid identities were denoted by total acyl carbon number and total double bond number.

Raw data from orbitrap mass spectrometers were processed using TraceFinder 3.3 software (Thermo Fisher Scientific; Waltham, MA) and Progenesis QI (Nonlinear Dynamics; Newcastle upon Tyne, UK) and targeted data from the QTRAP 5500 system were processed using MultiQuant (version 2.1, SCIEX; Framingham, MA). For each method, metabolite identities were confirmed using authentic reference standards or reference samples.

Thirty-nine individual metabolites belonging to four metabolite classes (11 LPC, 17 PC, 6 SM and 5 CER) were analyzed in this study. 12 were assessed with the HILIC-positive approach: C34:2 PC, C34:4 PC, C36:2 PC-A, C36:2 PC-B, C14:0 LPC, C22:5 LPC, C22:6 LPC, C14:0 SM, C16:0 SM, C18:0 SM, C20:0 SM, and C24:1 Ceramide. The remaining 27 metabolites were assessed with the C8-positive approach.

Supplementary Figures

**Supplementary Figure 1. Correlation among metabolites in premenopausal control samples at blood draw.** Pearson correlation was calculated for all pairs of individual metabolites, metabolite sums and metabolite ratios among premenopausal women at blood collection. Positive Pearson correlation coefficients are shown in shades of red and negative coefficients are shown in shades of blue.

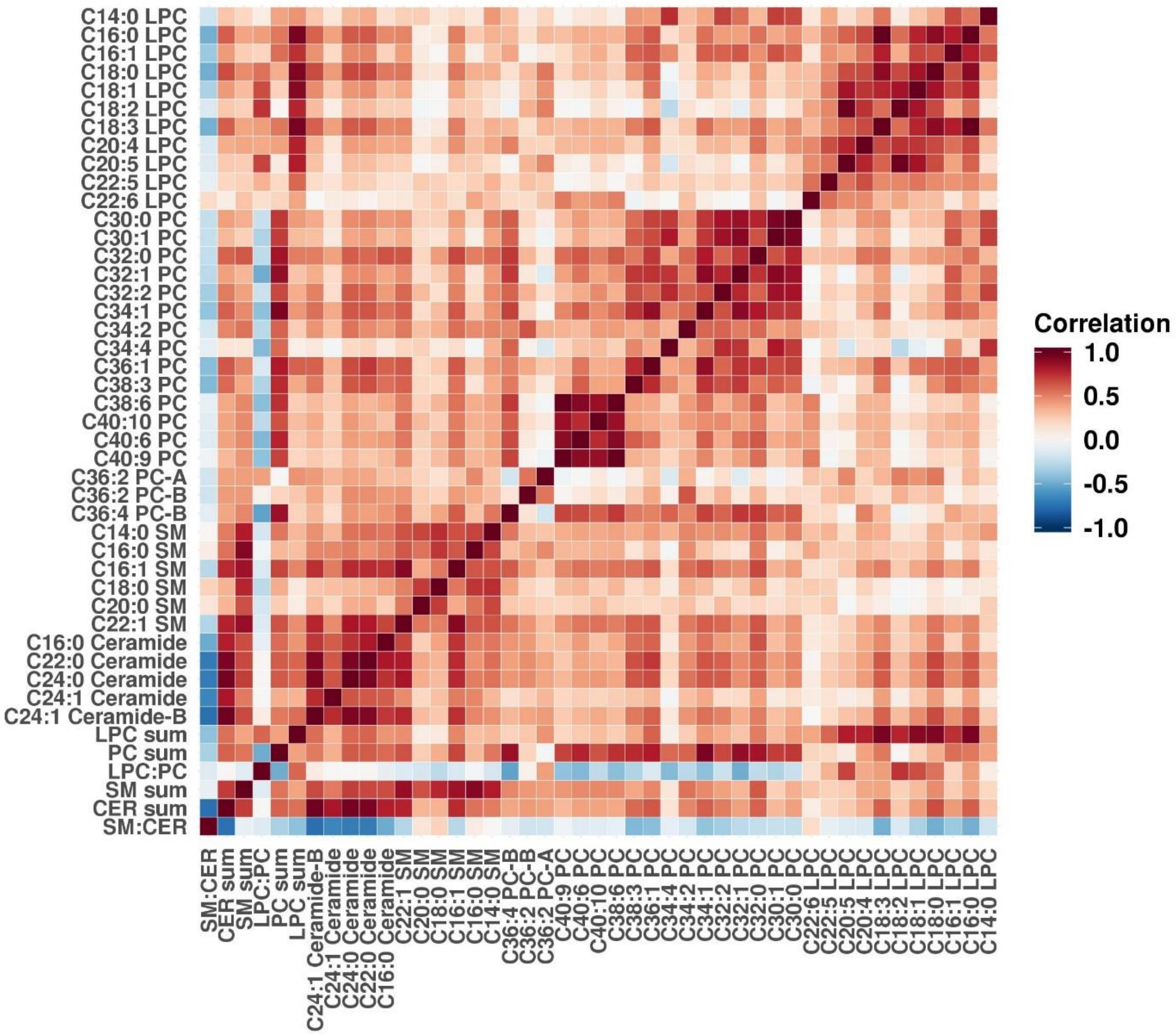

**Supplementary Figure 2. Correlation among metabolites in postmenopausal control samples at blood draw.** Pearson correlation was calculated for all pairs of individual metabolites, metabolite sums and metabolite ratios among postmenopausal women at blood collection. Positive Pearson correlation coefficients are shown in shades of red and negative coefficients are shown in shades of blue.

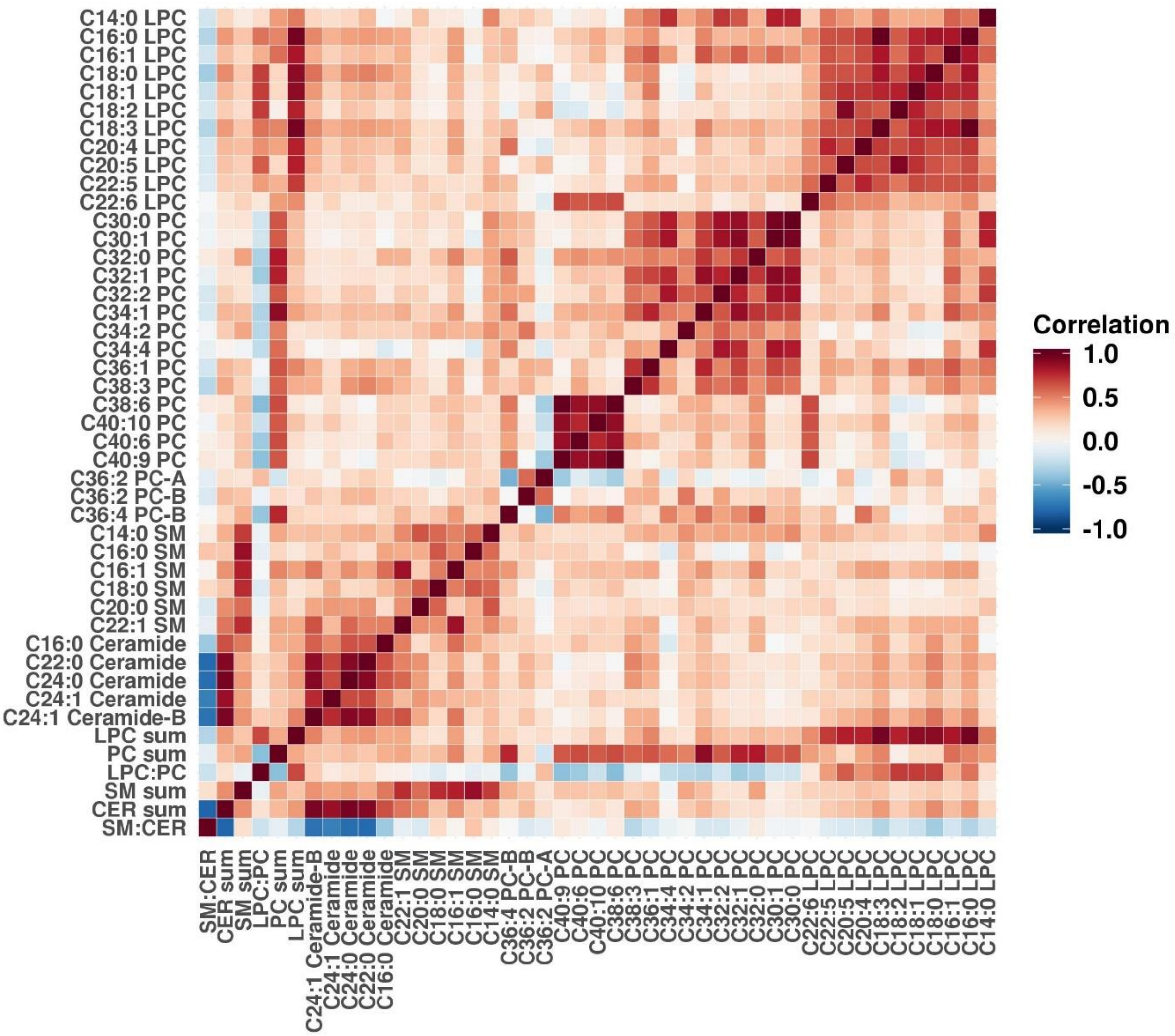

**Supplemental Figure 3 Odds ratios (OR) of risk of overall ovarian cancer by time between blood collection and diagnosis for an increase from the 10th to 90th percentile of metabolites levels.** OR>1 are shown in shades of red and OR<1 are shown in shades of blue. * p-value ≤0.05 (nominal significance), ** p-value ≤0.0125 (Bonferroni significance threshold).Estimates were adjusted for risk factors (duration of oral contraceptive use [none or <3 months, 3 months to 3 years, 3 years to 5 years, more than 5 years], tubal ligation [yes/no] and parity [no children, 1 child, 2 children, 3 children, 4+ children]) and additionally for matching factors (cohort [NHS, NHSII]; menopausal status and hormone therapy use at blood draw [premenopausal, postmenopausal and hormone therapy use, postmenopausal and no hormone therapy use, missing/unknown]; menopausal status at diagnosis [premenopausal, postmenopausal, or unknown]; age [±1 year], date of blood collection [± 1 month]; time of day of blood draw [±2 hours]; fasting status [>8 hours or ≤8 hours]) in analyses by time between blood collection and diagnosis.

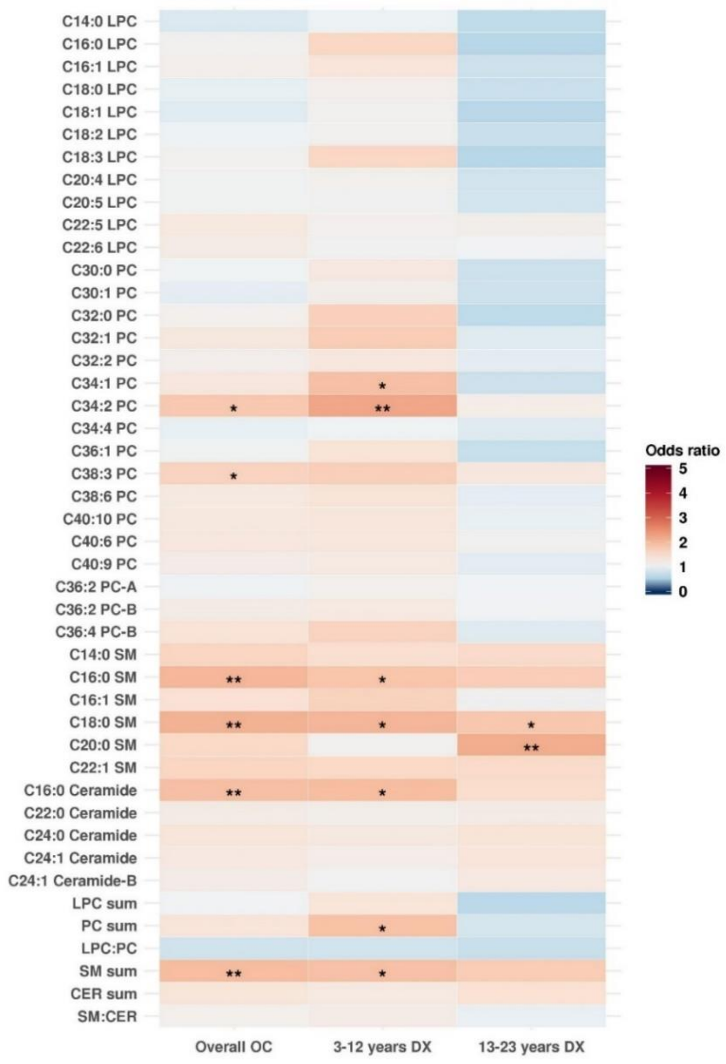

### Supplementary Tables

**Supplementary Table 1. Odds ratio (OR) for an increase from the 10th to 90th percentile of metabolite levels and 95% CI of ovarian cancer overall and according to histotype.**

| | | Overall OC | | Serous/PD OC | | Endo/CC OC | |
|---|---|---|---|---|---|---|---|
| **METABOLITE** | **HMDB_ID** | **PVAL** | **OR (95% CI)** | **PVAL** | **OR (95% CI)** | **PVAL** | **OR (95% CI)** |
| C14:0 LPC | HMDB10379 | 0.508 | 0.85 (0.54, 1.36) | 0.337 | 0.77 (0.46, 1.30) | 0.963 | 1.02 (0.39, 2.70) |
| C16:0 LPC | HMDB10382 | 0.758 | 1.08 (0.67, 1.75) | 0.651 | 0.89 (0.52, 1.50) | 0.683 | 0.82 (0.30, 2.17) |
| C16:1 LPC | HMDB10383* | 0.612 | 1.13 (0.71, 1.79) | 0.971 | 0.99 (0.60, 1.65) | 0.446 | 0.68 (0.25, 1.84) |
| C18:0 LPC | HMDB10384 | 0.851 | 0.95 (0.57, 1.58) | 0.957 | 0.98 (0.57, 1.71) | 0.041 | 0.32 (0.10, 0.93) |
| C18:1 LPC | HMDB02815* | 0.648 | 0.89 (0.55, 1.45) | 0.62 | 0.87 (0.51, 1.49) | 0.011 | 0.24 (0.07, 0.69) |
| C18:2 LPC | HMDB10386* | 0.928 | 0.98 (0.61, 1.57) | 0.805 | 0.93 (0.53, 1.62) | 0.015 | 0.26 (0.08, 0.75) |
| C18:3 LPC | HMDB10387* | 0.751 | 1.08 (0.67, 1.75) | 0.636 | 0.88 (0.52, 1.49) | 0.72 | 0.84 (0.31, 2.23) |
| C20:4 LPC | HMDB10395 | 0.85 | 1.05 (0.65, 1.70) | 0.427 | 0.81 (0.48, 1.36) | 0.862 | 0.92 (0.35, 2.41) |
| C20:5 LPC | HMDB10397 | 0.849 | 1.05 (0.65, 1.67) | 0.746 | 0.91 (0.53, 1.58) | 0.234 | 0.53 (0.18, 1.50) |
| C22:5 LPC | HMDB10403* | 0.266 | 1.30 (0.82, 2.08) | 0.857 | 1.05 (0.63, 1.76) | 0.89 | 1.07 (0.40, 2.89) |
| C22:6 LPC | HMDB10404 | 0.431 | 1.23 (0.73, 2.07) | 0.547 | 0.85 (0.50, 1.44) | 0.066 | 2.69 (0.96, 7.95) |
| C30:0 PC | HMDB07869* | 0.976 | 1.01 (0.63, 1.62) | 0.616 | 0.87 (0.52, 1.48) | 0.722 | 1.18 (0.48, 2.98) |
| C30:1 PC | HMDB07870* | 0.777 | 0.93 (0.58, 1.50) | 0.571 | 0.86 (0.51, 1.44) | 0.842 | 0.91 (0.36, 2.29) |
| C32:0 PC | HMDB07871* | 0.672 | 1.11 (0.68, 1.83) | 0.402 | 0.79 (0.46, 1.36) | 0.875 | 1.08 (0.40, 2.94) |
| C32:1 PC | HMDB07873* | 0.28 | 1.31 (0.80, 2.13) | 0.849 | 1.05 (0.62, 1.79) | 0.312 | 1.66 (0.63, 4.55) |
| C32:2 PC | HMDB07874* | 0.589 | 1.14 (0.71, 1.84) | 0.668 | 1.12 (0.66, 1.90) | 0.957 | 0.98 (0.39, 2.48) |
| C34:1 PC | HMDB07972* | 0.266 | 1.32 (0.81, 2.13) | 0.989 | 1.00 (0.59, 1.72) | 0.472 | 1.42 (0.54, 3.80) |
| C34:2 PC | HMDB07973* | 0.024 | 1.82 (1.08, 3.06) | 0.305 | 1.33 (0.77, 2.28) | 0.134 | 2.16 (0.80, 6.03) |
| C34:4 PC | HMDB07883* | 0.839 | 0.95 (0.59, 1.54) | 0.7 | 0.90 (0.53, 1.53) | 0.598 | 1.30 (0.49, 3.46) |
| C36:1 PC | HMDB08038* | 0.828 | 1.05 (0.67, 1.66) | 0.896 | 0.97 (0.58, 1.62) | 0.381 | 0.64 (0.24, 1.73) |
| C38:3 PC | HMDB08047* | 0.046 | 1.67 (1.01, 2.76) | 0.073 | 1.62 (0.96, 2.76) | 0.299 | 1.69 (0.63, 4.61) |
| C38:6 PC | HMDB07991* | 0.389 | 1.27 (0.73, 2.21) | 0.81 | 0.93 (0.53, 1.64) | 0.085 | 2.46 (0.90, 7.11) |
| C40:10 PC | HMDB08511* | 0.313 | 1.31 (0.78, 2.20) | 0.667 | 1.13 (0.65, 1.96) | 0.637 | 1.26 (0.49, 3.35) |
| C40:6 PC | HMDB08057* | 0.309 | 1.32 (0.77, 2.24) | 0.744 | 1.09 (0.64, 1.89) | 0.217 | 1.88 (0.70, 5.26) |
| C40:9 PC | HMDB08731* | 0.528 | 1.19 (0.69, 2.06) | 0.702 | 0.90 (0.51, 1.58) | 0.108 | 2.32 (0.85, 6.67) |
| C36:2 PC-A | HMDB08039* | 0.967 | 0.99 (0.61, 1.61) | 0.495 | 1.21 (0.70, 2.11) | 0.032 | 0.30 (0.09, 0.89) |
| C36:2 PC-B | HMDB08039* | 0.378 | 1.23 (0.77, 1.96) | 0.412 | 1.24 (0.74, 2.10) | 0.055 | 0.37 (0.13, 1.01) |

| | | | | | | | |
|---|---|---|---|---|---|---|---|
| C36:4 PC-B | HMDB08138* | 0.201 | 1.43 (0.83, 2.46) | 0.937 | 0.98 (0.56, 1.70) | 0.071 | 2.63 (0.93, 7.71) |
| C14:0 SM | HMDB12097 | 0.052 | 1.65 (1.00, 2.72) | 0.081 | 1.64 (0.94, 2.88) | 0.493 | 1.42 (0.52, 3.96) |
| C16:0 SM | HMDB10169 | 0.009 | 2.06 (1.19, 3.56) | 0.05 | 1.73 (1.00, 3.01) | 0.897 | 0.93 (0.33, 2.64) |
| C16:1 SM | NA | 0.166 | 1.44 (0.86, 2.41) | 0.426 | 1.25 (0.72, 2.17) | 0.976 | 1.02 (0.34, 2.99) |
| C18:0 SM | HMDB01348 | 0.004 | 2.10 (1.26, 3.49) | 0.056 | 1.67 (0.99, 2.84) | 0.011 | 4.06 (1.44, 12.60) |
| C20:0 SM | HMDB12102 | 0.063 | 1.60 (0.97, 2.63) | 0.031 | 1.80 (1.06, 3.08) | 0.676 | 1.24 (0.45, 3.36) |
| C22:1 SM | HMDB12104* | 0.059 | 1.65 (0.98, 2.77) | 0.108 | 1.58 (0.91, 2.75) | 0.727 | 0.82 (0.27, 2.44) |
| C16:0 Ceramide (d18:1) | HMDB04949 | 0.012 | 1.95 (1.16, 3.30) | 0.08 | 1.61 (0.95, 2.78) | 0.431 | 1.50 (0.54, 4.19) |
| C22:0 Ceramide (d18:1) | HMDB04952 | 0.442 | 1.21 (0.74, 1.97) | 0.502 | 1.20 (0.70, 2.05) | 0.633 | 1.28 (0.47, 3.52) |
| C24:0 Ceramide (d18:1) | HMDB04956 | 0.235 | 1.35 (0.82, 2.21) | 0.162 | 1.48 (0.86, 2.58) | 0.857 | 0.91 (0.32, 2.53) |
| C24:1 Ceramide (d18:1) | HMDB04953* | 0.304 | 1.29 (0.79, 2.11) | 0.412 | 1.24 (0.74, 2.11) | 0.314 | 1.70 (0.61, 4.85) |
| C24:1 Ceramide (d18:1)-B | HMDB04953* | 0.454 | 1.21 (0.74, 1.98) | 0.378 | 1.28 (0.74, 2.22) | 0.827 | 0.89 (0.31, 2.52) |
| LPC sum | NA | 0.928 | 1.02 (0.63, 1.65) | 0.655 | 0.89 (0.52, 1.51) | 0.145 | 0.47 (0.16, 1.28) |
| PC sum | NA | 0.21 | 1.40 (0.83, 2.37) | 0.978 | 1.01 (0.58, 1.75) | 0.181 | 1.99 (0.73, 5.59) |
| LPC:PC | NA | 0.3 | 0.76 (0.45, 1.28) | 0.556 | 0.84 (0.47, 1.50) | 0.011 | 0.24 (0.08, 0.71) |
| SM sum | NA | 0.011 | 1.97 (1.16, 3.32) | 0.045 | 1.77 (1.02, 3.10) | 0.83 | 1.12 (0.38, 3.28) |
| CER sum | NA | 0.209 | 1.37 (0.84, 2.24) | 0.186 | 1.44 (0.84, 2.49) | 0.662 | 1.26 (0.45, 3.54) |
| SM:CER | NA | 0.682 | 1.11 (0.68, 1.79) | 0.993 | 1.00 (0.59, 1.68) | 0.771 | 0.87 (0.33, 2.28) |

* representative HMDB ID

**Supplementary Table 2. Odds ratio (OR) for an increase from the 10th to 90th percentile of metabolite levels and 95% CI of ovarian cancer overall, for rapidly fatal and less aggressive tumors.**

| | | Overall OC | | Rapidly fatal | | Not-rapidly fatal | |
|---|---|---|---|---|---|---|---|
| **METABOLITE** | **HMDB_ID** | **PVAL** | **OR (95% CI)** | **PVAL** | **OR (95% CI)** | **PVAL** | **OR (95% CI)** |
| C14:0 LPC | HMDB10379 | 0.508 | 0.85 (0.54, 1.36) | 0.358 | 0.80 (0.50, 1.28) | 0.141 | 0.65 (0.37, 1.15) |
| C16:0 LPC | HMDB10382 | 0.758 | 1.08 (0.67, 1.75) | 0.923 | 1.02 (0.64, 1.65) | 0.346 | 0.75 (0.41, 1.36) |
| C16:1 LPC | HMDB10383* | 0.612 | 1.13 (0.71, 1.79) | 0.819 | 1.06 (0.66, 1.68) | 0.381 | 0.77 (0.43, 1.37) |
| C18:0 LPC | HMDB10384 | 0.851 | 0.95 (0.57, 1.58) | 0.825 | 0.95 (0.57, 1.56) | 0.455 | 0.79 (0.42, 1.47) |
| C18:1 LPC | HMDB02815* | 0.648 | 0.89 (0.55, 1.45) | 0.512 | 0.85 (0.52, 1.38) | 0.318 | 0.73 (0.40, 1.34) |
| C18:2 LPC | HMDB10386* | 0.928 | 0.98 (0.61, 1.57) | 0.647 | 0.89 (0.54, 1.47) | 0.667 | 0.87 (0.47, 1.62) |
| C18:3 LPC | HMDB10387* | 0.751 | 1.08 (0.67, 1.75) | 0.924 | 1.02 (0.64, 1.65) | 0.329 | 0.74 (0.40, 1.35) |
| C20:4 LPC | HMDB10395 | 0.85 | 1.05 (0.65, 1.70) | 0.945 | 0.98 (0.61, 1.58) | 0.422 | 0.79 (0.44, 1.40) |
| C20:5 LPC | HMDB10397 | 0.849 | 1.05 (0.65, 1.67) | 0.827 | 0.95 (0.57, 1.56) | 0.696 | 0.89 (0.48, 1.62) |
| C22:5 LPC | HMDB10403* | 0.266 | 1.30 (0.82, 2.08) | 0.589 | 1.14 (0.71, 1.83) | 0.875 | 1.05 (0.59, 1.87) |
| C22:6 LPC | HMDB10404 | 0.431 | 1.23 (0.73, 2.07) | 0.762 | 1.08 (0.67, 1.74) | 0.982 | 1.01 (0.56, 1.81) |
| C30:0 PC | HMDB07869* | 0.976 | 1.01 (0.63, 1.62) | 0.852 | 0.96 (0.60, 1.53) | 0.546 | 0.84 (0.48, 1.48) |
| C30:1 PC | HMDB07870* | 0.777 | 0.93 (0.58, 1.50) | 0.623 | 0.89 (0.56, 1.42) | 0.339 | 0.76 (0.43, 1.33) |
| C32:0 PC | HMDB07871* | 0.672 | 1.11 (0.68, 1.83) | 0.993 | 1.00 (0.62, 1.63) | 0.605 | 0.86 (0.47, 1.54) |
| C32:1 PC | HMDB07873* | 0.28 | 1.31 (0.80, 2.13) | 0.443 | 1.21 (0.75, 1.95) | 0.961 | 0.99 (0.55, 1.76) |
| C32:2 PC | HMDB07874* | 0.589 | 1.14 (0.71, 1.84) | 0.708 | 1.09 (0.69, 1.75) | 0.891 | 0.96 (0.54, 1.70) |
| C34:1 PC | HMDB07972* | 0.266 | 1.32 (0.81, 2.13) | 0.507 | 1.18 (0.73, 1.90) | 0.923 | 0.97 (0.54, 1.74) |
| C34:2 PC | HMDB07973* | 0.024 | 1.82 (1.08, 3.06) | 0.068 | 1.56 (0.97, 2.54) | 0.07 | 1.72 (0.96, 3.12) |
| C34:4 PC | HMDB07883* | 0.839 | 0.95 (0.59, 1.54) | 0.77 | 0.93 (0.58, 1.50) | 0.298 | 0.74 (0.41, 1.31) |
| C36:1 PC | HMDB08038* | 0.828 | 1.05 (0.67, 1.66) | 0.908 | 0.97 (0.61, 1.55) | 0.704 | 0.89 (0.50, 1.59) |
| C38:3 PC | HMDB08047* | 0.046 | 1.67 (1.01, 2.76) | 0.078 | 1.54 (0.96, 2.48) | 0.33 | 1.33 (0.75, 2.38) |
| C38:6 PC | HMDB07991* | 0.389 | 1.27 (0.73, 2.21) | 0.578 | 1.15 (0.70, 1.90) | 0.889 | 1.04 (0.57, 1.92) |
| C40:10 PC | HMDB08511* | 0.313 | 1.31 (0.78, 2.20) | 0.519 | 1.17 (0.72, 1.92) | 0.987 | 0.99 (0.55, 1.82) |
| C40:6 PC | HMDB08057* | 0.309 | 1.32 (0.77, 2.24) | 0.448 | 1.21 (0.74, 1.99) | 0.706 | 1.12 (0.62, 2.06) |
| C40:9 PC | HMDB08731* | 0.528 | 1.19 (0.69, 2.06) | 0.725 | 1.09 (0.66, 1.80) | 0.931 | 0.97 (0.53, 1.79) |
| C36:2 PC-A | HMDB08039* | 0.967 | 0.99 (0.61, 1.61) | 0.949 | 1.02 (0.62, 1.65) | 0.56 | 1.19 (0.66, 2.13) |
| C36:2 PC-B | HMDB08039* | 0.378 | 1.23 (0.77, 1.96) | 0.61 | 1.13 (0.71, 1.81) | 0.298 | 1.36 (0.76, 2.45) |
| C36:4 PC-B | HMDB08138* | 0.201 | 1.43 (0.83, 2.46) | 0.371 | 1.26 (0.76, 2.10) | 0.873 | 1.05 (0.58, 1.92) |
| C14:0 SM | HMDB12097 | 0.052 | 1.65 (1.00, 2.72) | 0.083 | 1.56 (0.95, 2.58) | 0.165 | 1.52 (0.84, 2.78) |

| | | | | | | | |
|---|---|---|---|---|---|---|---|
| C16:0 SM | HMDB10169 | 0.009 | 2.06 (1.19, 3.56) | 0.03 | 1.74 (1.06, 2.90) | 0.018 | 2.06 (1.14, 3.79) |
| C16:1 SM | NA | 0.166 | 1.44 (0.86, 2.41) | 0.244 | 1.35 (0.82, 2.23) | 0.477 | 1.25 (0.68, 2.29) |
| C18:0 SM | HMDB01348 | 0.004 | 2.10 (1.26, 3.49) | 0.008 | 1.91 (1.19, 3.09) | 0.011 | 2.10 (1.20, 3.75) |
| C20:0 SM | HMDB12102 | 0.063 | 1.60 (0.97, 2.63) | 0.061 | 1.57 (0.98, 2.53) | 0.132 | 1.54 (0.88, 2.73) |
| C22:1 SM | HMDB12104* | 0.059 | 1.65 (0.98, 2.77) | 0.067 | 1.59 (0.97, 2.63) | 0.29 | 1.39 (0.76, 2.57) |
| C16:0 Ceramide (d18:1) | HMDB04949 | 0.012 | 1.95 (1.16, 3.30) | 0.022 | 1.77 (1.09, 2.91) | 0.127 | 1.58 (0.88, 2.87) |
| C22:0 Ceramide (d18:1) | HMDB04952 | 0.442 | 1.21 (0.74, 1.97) | 0.355 | 1.26 (0.78, 2.04) | 0.598 | 1.17 (0.66, 2.09) |
| C24:0 Ceramide (d18:1) | HMDB04956 | 0.235 | 1.35 (0.82, 2.21) | 0.18 | 1.40 (0.86, 2.31) | 0.273 | 1.40 (0.77, 2.58) |
| C24:1 Ceramide (d18:1) | HMDB04953* | 0.304 | 1.29 (0.79, 2.11) | 0.31 | 1.28 (0.80, 2.06) | 0.515 | 1.21 (0.68, 2.15) |
| C24:1 Ceramide (d18:1)-B | HMDB04953* | 0.454 | 1.21 (0.74, 1.98) | 0.446 | 1.21 (0.74, 1.99) | 0.432 | 1.27 (0.70, 2.33) |
| LPC sum | NA | 0.928 | 1.02 (0.63, 1.65) | 0.86 | 0.96 (0.59, 1.56) | 0.333 | 0.74 (0.40, 1.36) |
| PC sum | NA | 0.21 | 1.40 (0.83, 2.37) | 0.403 | 1.24 (0.75, 2.05) | 0.98 | 1.01 (0.55, 1.85) |
| LPC:PC | NA | 0.3 | 0.76 (0.45, 1.28) | 0.328 | 0.77 (0.46, 1.30) | 0.272 | 0.70 (0.37, 1.32) |
| SM sum | NA | 0.011 | 1.97 (1.16, 3.32) | 0.02 | 1.83 (1.10, 3.05) | 0.04 | 1.88 (1.03, 3.48) |
| CER sum | NA | 0.209 | 1.37 (0.84, 2.24) | 0.165 | 1.42 (0.87, 2.33) | 0.283 | 1.39 (0.77, 2.53) |
| SM:CER | NA | 0.682 | 1.11 (0.68, 1.79) | 0.912 | 1.03 (0.64, 1.64) | 0.764 | 1.09 (0.62, 1.93) |

* representative HMDB ID

**Supplementary Table 3. Odds ratio (OR) for an increase from the 10th to 90th percentile of metabolite levels and 95% CI of ovarian cancer and by menopausal status at blood collection.**

| | | Overall OC | | Premenopausal | | Postmenopausal | |
|---|---|---|---|---|---|---|---|
| **METABOLITE** | **HMDB_ID** | **PVAL** | **OR (95% CI)** | **PVAL** | **OR (95% CI)** | **PVAL** | **OR (95% CI)** |
| C14:0 LPC | HMDB10379 | 0.508 | 0.85 (0.54, 1.36) | 0.747 | 0.87 (0.36, 2.08) | 0.327 | 0.72 (0.37, 1.39) |
| C16:0 LPC | HMDB10382 | 0.758 | 1.08 (0.67, 1.75) | 0.655 | 0.80 (0.30, 2.14) | 0.902 | 0.96 (0.50, 1.84) |
| C16:1 LPC | HMDB10383* | 0.612 | 1.13 (0.71, 1.79) | 0.37 | 0.64 (0.25, 1.68) | 0.636 | 1.16 (0.63, 2.14) |
| C18:0 LPC | HMDB10384 | 0.851 | 0.95 (0.57, 1.58) | 0.346 | 0.61 (0.22, 1.71) | 0.724 | 0.88 (0.44, 1.76) |
| C18:1 LPC | HMDB02815* | 0.648 | 0.89 (0.55, 1.45) | 0.2 | 0.55 (0.22, 1.38) | 0.998 | 1.00 (0.52, 1.91) |
| C18:2 LPC | HMDB10386* | 0.928 | 0.98 (0.61, 1.57) | 0.568 | 0.80 (0.37, 1.72) | 0.929 | 1.03 (0.52, 2.04) |
| C18:3 LPC | HMDB10387* | 0.751 | 1.08 (0.67, 1.75) | 0.635 | 0.79 (0.30, 2.10) | 0.911 | 0.96 (0.50, 1.85) |
| C20:4 LPC | HMDB10395 | 0.85 | 1.05 (0.65, 1.70) | 0.132 | 0.47 (0.17, 1.26) | 0.818 | 1.08 (0.55, 2.14) |
| C20:5 LPC | HMDB10397 | 0.849 | 1.05 (0.65, 1.67) | 0.824 | 0.92 (0.43, 1.98) | 0.86 | 0.94 (0.48, 1.86) |
| C22:5 LPC | HMDB10403* | 0.266 | 1.30 (0.82, 2.08) | 0.275 | 0.62 (0.26, 1.47) | 0.209 | 1.55 (0.78, 3.08) |
| C22:6 LPC | HMDB10404 | 0.431 | 1.23 (0.73, 2.07) | 0.255 | 1.74 (0.67, 4.55) | 0.777 | 0.90 (0.44, 1.84) |
| C30:0 PC | HMDB07869* | 0.976 | 1.01 (0.63, 1.62) | 0.736 | 0.85 (0.33, 2.20) | 0.745 | 0.90 (0.47, 1.71) |
| C30:1 PC | HMDB07870* | 0.777 | 0.93 (0.58, 1.50) | 0.566 | 0.76 (0.30, 1.94) | 0.492 | 0.79 (0.41, 1.54) |
| C32:0 PC | HMDB07871* | 0.672 | 1.11 (0.68, 1.83) | 0.413 | 0.68 (0.27, 1.70) | 0.555 | 1.23 (0.62, 2.42) |
| C32:1 PC | HMDB07873* | 0.28 | 1.31 (0.80, 2.13) | 0.88 | 1.07 (0.44, 2.63) | 0.78 | 1.10 (0.56, 2.19) |
| C32:2 PC | HMDB07874* | 0.589 | 1.14 (0.71, 1.84) | 0.679 | 0.81 (0.31, 2.16) | 0.561 | 1.22 (0.63, 2.35) |
| C34:1 PC | HMDB07972* | 0.266 | 1.32 (0.81, 2.13) | 0.861 | 0.92 (0.39, 2.21) | 0.651 | 1.17 (0.59, 2.34) |
| C34:2 PC | HMDB07973* | 0.024 | 1.82 (1.08, 3.06) | 0.682 | 1.22 (0.47, 3.16) | 0.031 | 2.29 (1.08, 4.86) |
| C34:4 PC | HMDB07883* | 0.839 | 0.95 (0.59, 1.54) | 0.299 | 0.61 (0.24, 1.55) | 0.936 | 0.97 (0.49, 1.95) |
| C36:1 PC | HMDB08038* | 0.828 | 1.05 (0.67, 1.66) | 0.366 | 0.65 (0.25, 1.66) | 0.853 | 0.94 (0.50, 1.79) |
| C38:3 PC | HMDB08047* | 0.046 | 1.67 (1.01, 2.76) | 0.93 | 1.05 (0.39, 2.81) | 0.364 | 1.40 (0.67, 2.92) |
| C38:6 PC | HMDB07991* | 0.389 | 1.27 (0.73, 2.21) | 0.594 | 1.29 (0.50, 3.33) | 0.506 | 1.30 (0.60, 2.78) |
| C40:10 PC | HMDB08511* | 0.313 | 1.31 (0.78, 2.20) | 0.764 | 1.16 (0.45, 2.97) | 0.282 | 1.46 (0.73, 2.91) |
| C40:6 PC | HMDB08057* | 0.309 | 1.32 (0.77, 2.24) | 0.621 | 1.26 (0.50, 3.21) | 0.59 | 1.23 (0.58, 2.59) |
| C40:9 PC | HMDB08731* | 0.528 | 1.19 (0.69, 2.06) | 0.657 | 1.24 (0.48, 3.15) | 0.607 | 1.22 (0.57, 2.60) |
| C36:2 PC-A | HMDB08039* | 0.967 | 0.99 (0.61, 1.61) | 0.635 | 0.77 (0.26, 2.26) | 0.811 | 1.09 (0.55, 2.12) |
| C36:2 PC-B | HMDB08039* | 0.378 | 1.23 (0.77, 1.96) | 0.773 | 0.86 (0.30, 2.44) | 0.505 | 1.25 (0.65, 2.40) |
| C36:4 PC-B | HMDB08138* | 0.201 | 1.43 (0.83, 2.46) | 0.278 | 0.56 (0.20, 1.59) | 0.152 | 1.76 (0.81, 3.79) |
| C14:0 SM | HMDB12097 | 0.052 | 1.65 (1.00, 2.72) | 0.379 | 0.62 (0.22, 1.78) | 0.02 | 2.24 (1.13, 4.41) |

| | | | | | | | |
|---|---|---|---|---|---|---|---|
| C16:0 SM | HMDB10169 | 0.009 | 2.06 (1.19, 3.56) | 0.613 | 0.77 (0.27, 2.15) | 0.006 | 2.98 (1.37, 6.51) |
| C16:1 SM | NA | 0.166 | 1.44 (0.86, 2.41) | 0.125 | 0.44 (0.16, 1.25) | 0.017 | 2.50 (1.18, 5.29) |
| C18:0 SM | HMDB01348 | 0.004 | 2.10 (1.26, 3.49) | 0.097 | 2.49 (0.85, 7.29) | 0.041 | 2.02 (1.03, 3.94) |
| C20:0 SM | HMDB12102 | 0.063 | 1.60 (0.97, 2.63) | 0.738 | 0.85 (0.32, 2.22) | 0.06 | 1.86 (0.97, 3.56) |
| C22:1 SM | HMDB12104* | 0.059 | 1.65 (0.98, 2.77) | 0.069 | 0.36 (0.12, 1.08) | 0.003 | 3.11 (1.46, 6.62) |
| C16:0 Ceramide (d18:1) | HMDB04949 | 0.012 | 1.95 (1.16, 3.30) | 0.285 | 1.76 (0.62, 4.99) | 0.039 | 2.16 (1.04, 4.48) |
| C22:0 Ceramide (d18:1) | HMDB04952 | 0.442 | 1.21 (0.74, 1.97) | 0.665 | 0.80 (0.30, 2.17) | 0.549 | 1.22 (0.63, 2.37) |
| C24:0 Ceramide (d18:1) | HMDB04956 | 0.235 | 1.35 (0.82, 2.21) | 0.639 | 0.79 (0.30, 2.11) | 0.46 | 1.28 (0.66, 2.49) |
| C24:1 Ceramide (d18:1) | HMDB04953* | 0.304 | 1.29 (0.79, 2.11) | 0.416 | 1.48 (0.58, 3.79) | 0.819 | 1.08 (0.55, 2.12) |
| C24:1 Ceramide (d18:1)-B | HMDB04953* | 0.454 | 1.21 (0.74, 1.98) | 0.455 | 0.68 (0.25, 1.88) | 0.479 | 1.27 (0.66, 2.44) |
| LPC sum | NA | 0.928 | 1.02 (0.63, 1.65) | 0.369 | 0.65 (0.26, 1.65) | 0.999 | 1.00 (0.52, 1.93) |
| PC sum | NA | 0.21 | 1.40 (0.83, 2.37) | 0.635 | 0.80 (0.31, 2.02) | 0.377 | 1.40 (0.66, 2.98) |
| LPC:PC | NA | 0.3 | 0.76 (0.45, 1.28) | 0.587 | 0.78 (0.32, 1.91) | 0.502 | 0.78 (0.37, 1.63) |
| SM sum | NA | 0.011 | 1.97 (1.16, 3.32) | 0.352 | 0.61 (0.22, 1.72) | 0.003 | 3.22 (1.51, 6.86) |
| CER sum | NA | 0.209 | 1.37 (0.84, 2.24) | 0.988 | 1.01 (0.38, 2.70) | 0.469 | 1.28 (0.66, 2.51) |
| SM:CER | NA | 0.682 | 1.11 (0.68, 1.79) | 0.42 | 0.68 (0.26, 1.75) | 0.151 | 1.64 (0.84, 3.22) |
| * representative HMDB ID | | | | | | | |

**Supplementary Table 4. Odds ratio (OR) for an increase from the 10th to 90th percentile of metabolite levels and 95% CI of ovarian cancer and by time between blood collection and diagnosis.**

| | | Overall OC | | 3-12 years to DX | | 13-23 years to DX | |
|---|---|---|---|---|---|---|---|
| **METABOLITE** | **HMDB_ID** | **PVAL** | **OR (95% CI)** | **PVAL** | **OR (95% CI)** | **PVAL** | **OR (95% CI)** |
| C14:0 LPC | HMDB10379 | 0.508 | 0.85 (0.54, 1.36) | 0.958 | 0.98 (0.54, 1.79) | 0.109 | 0.63 (0.35, 1.10) |
| C16:0 LPC | HMDB10382 | 0.758 | 1.08 (0.67, 1.75) | 0.121 | 1.63 (0.88, 3.03) | 0.06 | 0.57 (0.32, 1.02) |
| C16:1 LPC | HMDB10383* | 0.612 | 1.13 (0.71, 1.79) | 0.285 | 1.38 (0.77, 2.49) | 0.294 | 0.74 (0.42, 1.30) |
| C18:0 LPC | HMDB10384 | 0.851 | 0.95 (0.57, 1.58) | 0.691 | 1.14 (0.60, 2.18) | 0.265 | 0.71 (0.39, 1.29) |
| C18:1 LPC | HMDB02815* | 0.648 | 0.89 (0.55, 1.45) | 0.777 | 1.09 (0.59, 2.03) | 0.08 | 0.58 (0.32, 1.06) |
| C18:2 LPC | HMDB10386* | 0.928 | 0.98 (0.61, 1.57) | 0.804 | 1.09 (0.57, 2.08) | 0.26 | 0.70 (0.37, 1.30) |
| C18:3 LPC | HMDB10387* | 0.751 | 1.08 (0.67, 1.75) | 0.118 | 1.63 (0.89, 3.04) | 0.057 | 0.57 (0.31, 1.01) |
| C20:4 LPC | HMDB10395 | 0.85 | 1.05 (0.65, 1.70) | 0.709 | 1.12 (0.62, 2.03) | 0.362 | 0.77 (0.43, 1.36) |
| C20:5 LPC | HMDB10397 | 0.849 | 1.05 (0.65, 1.67) | 0.84 | 1.07 (0.56, 2.03) | 0.427 | 0.78 (0.42, 1.44) |
| C22:5 LPC | HMDB10403* | 0.266 | 1.30 (0.82, 2.08) | 0.758 | 1.10 (0.60, 2.02) | 0.61 | 1.16 (0.66, 2.03) |
| C22:6 LPC | HMDB10404 | 0.431 | 1.23 (0.73, 2.07) | 0.822 | 1.07 (0.58, 2.00) | 0.917 | 1.03 (0.58, 1.85) |
| C30:0 PC | HMDB07869* | 0.976 | 1.01 (0.63, 1.62) | 0.384 | 1.29 (0.73, 2.31) | 0.313 | 0.74 (0.41, 1.32) |
| C30:1 PC | HMDB07870* | 0.777 | 0.93 (0.58, 1.50) | 0.623 | 1.16 (0.65, 2.06) | 0.28 | 0.73 (0.41, 1.29) |
| C32:0 PC | HMDB07871* | 0.672 | 1.11 (0.68, 1.83) | 0.092 | 1.70 (0.92, 3.19) | 0.133 | 0.63 (0.34, 1.15) |
| C32:1 PC | HMDB07873* | 0.28 | 1.31 (0.80, 2.13) | 0.065 | 1.77 (0.97, 3.27) | 0.661 | 0.88 (0.48, 1.58) |
| C32:2 PC | HMDB07874* | 0.589 | 1.14 (0.71, 1.84) | 0.36 | 1.32 (0.73, 2.38) | 0.743 | 0.91 (0.51, 1.61) |
| C34:1 PC | HMDB07972* | 0.266 | 1.32 (0.81, 2.13) | 0.034 | 1.94 (1.05, 3.60) | 0.325 | 0.74 (0.40, 1.34) |
| C34:2 PC | HMDB07973* | 0.024 | 1.82 (1.08, 3.06) | 0.012 | 2.24 (1.20, 4.26) | 0.559 | 1.18 (0.67, 2.10) |
| C34:4 PC | HMDB07883* | 0.839 | 0.95 (0.59, 1.54) | 0.986 | 1.01 (0.55, 1.83) | 0.646 | 0.87 (0.49, 1.56) |
| C36:1 PC | HMDB08038* | 0.828 | 1.05 (0.67, 1.66) | 0.249 | 1.42 (0.78, 2.58) | 0.226 | 0.69 (0.38, 1.25) |
| C38:3 PC | HMDB08047* | 0.046 | 1.67 (1.01, 2.76) | 0.07 | 1.73 (0.96, 3.16) | 0.35 | 1.32 (0.74, 2.38) |
| C38:6 PC | HMDB07991* | 0.389 | 1.27 (0.73, 2.21) | 0.294 | 1.41 (0.75, 2.69) | 0.786 | 0.92 (0.51, 1.67) |
| C40:10 PC | HMDB08511* | 0.313 | 1.31 (0.78, 2.20) | 0.331 | 1.36 (0.73, 2.54) | 0.869 | 0.95 (0.53, 1.71) |
| C40:6 PC | HMDB08057* | 0.309 | 1.32 (0.77, 2.24) | 0.36 | 1.34 (0.72, 2.51) | 0.775 | 1.09 (0.60, 1.98) |
| C40:9 PC | HMDB08731* | 0.528 | 1.19 (0.69, 2.06) | 0.473 | 1.26 (0.67, 2.40) | 0.769 | 0.91 (0.50, 1.66) |
| C36:2 PC-A | HMDB08039* | 0.967 | 0.99 (0.61, 1.61) | 0.746 | 1.11 (0.59, 2.10) | 0.988 | 1.00 (0.55, 1.85) |
| C36:2 PC-B | HMDB08039* | 0.378 | 1.23 (0.77, 1.96) | 0.453 | 1.26 (0.69, 2.30) | 0.948 | 1.02 (0.57, 1.84) |
| C36:4 PC-B | HMDB08138* | 0.201 | 1.43 (0.83, 2.46) | 0.112 | 1.67 (0.89, 3.14) | 0.688 | 0.88 (0.47, 1.64) |
| C14:0 SM | HMDB12097 | 0.052 | 1.65 (1.00, 2.72) | 0.192 | 1.51 (0.82, 2.81) | 0.141 | 1.58 (0.86, 2.93) |

| | | | | | | | |
|---|---|---|---|---|---|---|---|
| C16:0 SM | HMDB10169 | 0.009 | 2.06 (1.19, 3.56) | 0.048 | 1.87 (1.01, 3.50) | 0.075 | 1.75 (0.95, 3.27) |
| C16:1 SM | NA | 0.166 | 1.44 (0.86, 2.41) | 0.102 | 1.69 (0.90, 3.20) | 0.818 | 1.07 (0.58, 1.98) |
| C18:0 SM | HMDB01348 | 0.004 | 2.10 (1.26, 3.49) | 0.019 | 2.07 (1.13, 3.85) | 0.035 | 1.84 (1.05, 3.28) |
| C20:0 SM | HMDB12102 | 0.063 | 1.60 (0.97, 2.63) | 0.773 | 1.09 (0.60, 2.00) | 0.009 | 2.18 (1.22, 3.94) |
| C22:1 SM | HMDB12104* | 0.059 | 1.65 (0.98, 2.77) | 0.146 | 1.60 (0.85, 3.02) | 0.151 | 1.57 (0.85, 2.92) |
| C16:0 Ceramide (d18:1) | HMDB04949 | 0.012 | 1.95 (1.16, 3.30) | 0.035 | 1.96 (1.06, 3.67) | 0.145 | 1.54 (0.86, 2.78) |
| C22:0 Ceramide (d18:1) | HMDB04952 | 0.442 | 1.21 (0.74, 1.97) | 0.634 | 1.16 (0.63, 2.16) | 0.497 | 1.22 (0.69, 2.18) |
| C24:0 Ceramide (d18:1) | HMDB04956 | 0.235 | 1.35 (0.82, 2.21) | 0.448 | 1.27 (0.68, 2.38) | 0.254 | 1.42 (0.78, 2.59) |
| C24:1 Ceramide (d18:1) | HMDB04953* | 0.304 | 1.29 (0.79, 2.11) | 0.605 | 1.17 (0.65, 2.12) | 0.276 | 1.38 (0.78, 2.45) |
| C24:1 Ceramide (d18:1)-B | HMDB04953* | 0.454 | 1.21 (0.74, 1.98) | 0.86 | 1.06 (0.57, 1.98) | 0.4 | 1.29 (0.71, 2.35) |
| LPC sum | NA | 0.928 | 1.02 (0.63, 1.65) | 0.298 | 1.39 (0.75, 2.61) | 0.081 | 0.59 (0.32, 1.06) |
| PC sum | NA | 0.21 | 1.40 (0.83, 2.37) | 0.044 | 1.91 (1.02, 3.60) | 0.467 | 0.80 (0.43, 1.47) |
| LPC:PC | NA | 0.3 | 0.76 (0.45, 1.28) | 0.447 | 0.77 (0.39, 1.51) | 0.27 | 0.70 (0.37, 1.32) |
| SM sum | NA | 0.011 | 1.97 (1.16, 3.32) | 0.04 | 1.95 (1.04, 3.71) | 0.069 | 1.77 (0.96, 3.31) |
| CER sum | NA | 0.209 | 1.37 (0.84, 2.24) | 0.461 | 1.26 (0.68, 2.33) | 0.204 | 1.47 (0.81, 2.67) |
| SM:CER | NA | 0.682 | 1.11 (0.68, 1.79) | 0.515 | 1.22 (0.67, 2.23) | 0.885 | 0.96 (0.55, 1.69) |
| * representative HMDB ID | | | | | | | |


1. Tworoger SS, Sluss P, Hankinson SE. Association between plasma prolactin concentrations and risk of breast cancer among predominately premenopausal women. Cancer research. 2006;66(4):2476-82.
